# Efficient Derivation of Immortalized, Isogenic Cell Lines from Genetically Defined Murine Hepatoblastomas

**DOI:** 10.1101/2024.09.29.615692

**Authors:** Huabo Wang, Jie Lu, Keyao Chen, Bingwei Ma, Colin Henchy, Jessica Knapp, Sarangarajan Ranganathan, Edward V. Prochownik

## Abstract

**Background & Aims:** Molecularly, hepatoblastoma (HB), the most common childhood liver cancer, is the simplest of all human neoplasms, with the vast majority deregulating the Wnt/β-catenin, Hippo and/or NFE2/NRF2 signaling pathways. Murine HBs can be generated by over-expressing any pairwise or triple combination of mutant forms of these pathways’ terminal effectors, namely β-catenin (B), YAP (Y) and NFE2L2/NRF (N). Each molecular subtypes displays distinct features resembling those of human HBs. However, research has been hampered by a paucity of established cell lines of any species.

**Methods:** We show here that immortalized cell lines can be routinely established from murine HBs that over-express B+Y and B+Y+N. This is facilitated by the concurrent *in vivo,* Crispr-mediated inactivation of the *Cdkn2a* tumor suppressor locus.

**Results:** Eight BY and 3 BYN cell lines have been generated and characterized and are available to the HB research community. Ten of these lines can be regrown as subcutaneous and metastatic lung tumors in the immuno-competent mice from which they originated while retaining their original histologic features. During maintenance as spheroids *in vitro*, or during *in vivo* propagation, tumor cells express endothelial cell markers, particularly in regions that are hypoxic and/or in proximity to incipient blood vessels.

**Conclusions:** The ability to generate isogenic HB cell lines with defined oncogenic drivers should facilitate studies that are best performed *in vitro*. The approach may also be useful for deriving HB cell lines associated with less common molecular drivers and from human tumors.

**Synopsis:** The derivation of multiple immortalized murine hepatoblastoma cell lines driven by defined oncogenes is described. These lines are isogenic, retain their tumorigenicity in immuno-competent mice, readily form spheroids and express endothelial markers in response to hypoxia. They will allow studies that have heretofore been difficult or impossible to perform *in vivo*.

## INTRODUCTION

Hepatoblastoma (HB) is the most common pediatric liver cancer, with nearly all cases occurring prior to the age of 4 years. ^1–6^ At least 7 distinct histo-pathologic subtypes are known and, along with transcriptional profiling, are useful for identifying and stratifying high-risk individuals. ^3–10^ While surgery and chemotherapy are curative in about 70-80% of cases, the remaining ones will either progress or recur. Unfortunately, cures in these latter groups can only be achieved with orthotopic liver transplantation, which in many cases is cost-prohibitive, of limited accessibility and associated with significant long-term morbidities in its own right ^11^. For this reason, and because the world-wide incidence of HB is increasing faster than that of any other pediatric cancer, there exists an immediate need to identify new and more effective therapeutic options. ^12, 13^

The molecular underpinnings of HB first emerged following reports that mutations of the transcription factor (TF) β-catenin (B), the terminal effector of the Wnt/β-catenin pathway, could be detected in up to 80% of human tumors. ^1, 14–17^ Deregulation of the Hippo tumor suppressor pathway was subsequently described, which led to the demonstration that over-expressing a constitutively active form of the Hippo terminal effector TF “<u>y</u>es-<u>a</u>ssociated <u>p</u>rotein” (YAP^S127A^) (Y), together with a patient-derived B mutant (B^del90^), could induce liver tumors bearing a close resemblance human HBs. ^18–21^ A third molecular abnormality involving amplification or mutation of the gene encoding the oxidative stress response TF NFE2L2/NRF2 (N) was subsequently identified in over half of HBs ^20^. Thus, deregulation and/or mutation of these 3 TFs in various combinations accounts for the vast majority of the molecular abnormalities associated with HB, which otherwise bears few other common mutations and is the least mutationally complex of all human cancers. ^22, 23^ We subsequently demonstrated that any pairwise combination of the above B, Y and N mutants, delivered in Sleeping Beauty (SB) vectors directly to the livers of mice, could efficiently induce HBs and that the 3 together generated tumors with particularly aggressive behaviors. ^20^ Indeed, each B,Y and N combination (BY, NY, BN and BYN) generated HBs with distinct growth rates, histologies, metabolic and biochemical features and transcriptional profiles that often mimicked those of human HBs. ^20, 21^ We also demonstrated that BY HB behaviors were further influenced by the identity of the B mutation. ^21^ That even WT B could induce HBs in association with Y was consistent with the fact that human HBs occasionally harbor germ-line or acquired inactivating mutations in the *APC* gene that allow for the constitutive localization of WT B to the nucleus. *APC* encodes an integral component of the APC tumor suppressor (TS) complex that normally maintains B in a highly degradable cytoplasmic form that is only released, stabilized and translocated to the nucleus in response to Wnt pathway signaling. ^24, 25^ Germ-line *APC* mutations and permanent nuclear residency of WT B are the basis of the familial adenomatous polyposis (FAP) syndrome, which, in addition to its association with a near 100% life-time incidence of colo-rectal cancer, contributes to a several thousand-fold increase in HB in early childhood. ^14, 24, 26, 27^ Rare cases of HB harboring acquired mutations in genes encoding other components of the APC complex such as Axin 1 and Axin 2 are associated with a similar stabilization and constitutive nuclear localization of B. ^17^

A long-standing and serious impediment to HB research is the acknowledged paucity of established human or murine HB cell lines of any type, least of all those that can be propagated in immuno-competent mice and/or that reflect the tumor’s authentic drivers and molecular heterogeneity. ^28^ Such lines would potentially facilitate or simplify studies that are ideally performed *in vitro* such as those aimed at identifying novel therapeutic agents or genetic vulnerabilities or requiring strict control over the extracellular environment. ^29^ Because the various murine HBs described above possess very different biological, biochemical, metabolic and transcriptional profiles ^20, 21^ cell lines with defined oncogenic drivers would also allow for carefully controlled comparative studies. Unfortunately, only a handful of HB cell lines currently exist, with the most widely used HepG2 human line being the only one available from the American Type Culture Collection (ATCC). ^30–32^ Reported in 1979 and established from the tumor of a 15 year old boy, this cell line was initially, and in some cases continues to be, mis-represented as originating from a hepatocellular carcinoma (HCC) although its HB derivation was subsequently verified. ^30, 33, 34^ Even so, the advanced age of the patient raises the question as to how representative HepG2 cells are of typical HBs as does the fact that, despite their containing nearly 100 different mutations, none involve B, Y or N or even some of the less common HB-associated driver genes (https://www.cccells.org/tables/Hepatobalstoma.php). The cells have also been subjected to considerable *in vitro* adaptation, genetic drift and sub-line selection since their derivation. Recently described murine HB cell lines are alleged to resemble aggressive human HBs but are driven by Myc rather than by any of the B,Y and N combinations that underlie nearly all human HBs. The cytoplasmic localization of B in these tumors is also not representative of its behavior in the majority of human cases. ^1, 14, 21, 29^ Moreover, liver cancers driven solely by Myc have been previously described as being more representative of undifferentiated HCCs ^35^. Finally, B+Y can still drive HB tumorigenesis in mice in Myc’s absence, thus showing it to be dispensable for tumor initiation, although it does contribute to tumor grow rate. ^20^ Thus Myc-driven murine tumors are not molecularly representative of naturally occurring human or other experimental murine HBs. ^1, 5, 29^

The human *CDKN2A* gene encodes 2 TSs: p16^INK4A^ and p14^ARF^ (p19^ARF^ in mice). Initiating from different promoters and first exons (1α and 1β, respectively), Cdkn2a transcripts share common second exon sequences that are translated in alternate reading frames. ^36, 37^ Indirectly responsible for maintaining transcriptionally inert complexes between the retinoblastoma (Rb) protein and members of the E2F TF family via its inhibition of cyclin-dependent kinases 4 and 6 (Cdk4/6), p16^INK4A^ contributes to the onset of cell cycle arrest and senescence. ^38–40^ By virtue of inhibiting the Mdm2 E3 ubiquitin ligase, p19^ARF^ indirectly stabilizes and activates TP53 so as to promote cell cycle arrest and apoptosis, particularly in response to oncogenic or genotoxic insults. ^37, 40, 41^ Although we have shown that the enforced co-expression of p19^ARF^ during the generation of BY HBs completely blocks tumorigenesis and others have demonstrated that hypermethylation and/or down-regulation of the *CDKN2A* locus occurs in a subset of human HBs, wider roles for p16^INK4A^ and p19^ARF^ in the pathogenesis of HB have not been reported. ^42–44^ However, inactivation of p16*^INK4A^*and/or p19^ARF^ is necessary to achieve immortalization of numerous human and murine cell types by allowing them to escape the proliferative barriers imposed by replicative and/or oncogence-mediated senescent and/or apoptotic signals. ^36, 41, 45–50^ The *CDKN2A* locus is also commonly mutated, deleted or epigenetically inactivated in HBs and a variety of human and murine cancers. ^39, 40, 42, 44, 51^

Using the above observations as a guide, we show here that *in vivo* Crispr/Cas9-mediated targeting of *Cdkn2a* exon 2 or the more selective targeting of p16^INK4a^-encoding exon 1α is sufficient to generate immortalized BY or BYN cell lines from primary tumors. The 11 such cell lines that have been derived from molecularly defined and otherwise isogenic HBs will allow for a variety of comparative studies that have heretofore been difficult or impossible to conduct *in vivo*.

## METHODS

### Animal care and husbandry

All animal care, husbandry and procedures were performed in compliance with the Public Health Service Policy on Humane Care and Use of Laboratory Animal Research Guide for Care and Use of Laboratory Animals. All experimental procedures, diets and tests were approved by the Institutional Animal Care and Use Committee (IACUC) at the University of Pittsburgh. Mice were maintained in a specific pathogen-free facility at UPMC Children’s Hospital of Pittsburgh, housed in micro-isolator cages and provided with water and standard animal chow *ad libitum*.

### Plasmids and transfections

WT p16^INK4A^- and p19^ARF^-encoding cDNAs were amplified by RT-PCR from HB samples using a SuperScript IV 1^st^ Strand cDNA Synthesis Kit (Life Technologies/Thermo Fisher, Inc., Waltham, MA) and Pfu DNA polymerase (Promega, Inc. Madison, WI) according to the directions of the respective vendors. The PCR primers used to amplify each of these are shown in Table 1. cDNAs were then cloned into the pSBbi-RP Sleeping Beauty (SB) expression vector that also encodes dTomato and puromycin resistance (Addgene, Inc., Waltham, MA). After purification using DNeasy columns (Qiagen, Inc. Germantown, MD), the plasmids were transfected into HB cell lines in the presence of a 10-fold lower molar amount of a pCMV-SB transposase-encoding vector using Lipofectamine (Thermo-Fisher). ^20, 21^ Expression was confirmed 2 days later by western blotting for the respective protein of interest and for the presence of dTomato by fluorescence microscopy. The subsequent proliferation of these cells was then monitored by re-plating the transfected population into 12 well dishes at low density in the presence of puromycin (2 μg/ml) and monitoring the subsequent expansion of dTomato+ cells on an Incucyte SX5 live cell imaging instrument (Sartorius Instruments, Göttingen, Germany). For other experiments involving the expression of p16^INK4A^- p19^ARF^ fusion proteins, the pSBbi-RP vector was first modified so as to encode a V5 epitope (pSBbi-RP-V5) that could be added to the C-terminus of any expressed protein. cDNAs encoding the desired p16^INK4A^-p19^ARF^ fusion or mutant proteins were then cloned upstream of this site to allow them to be expressed as V5 epitope-tagged fusion proteins, which were detected with an anti-V5 antibody (ThermoFisher, Inc., Waltham MA).

**Table 1.**
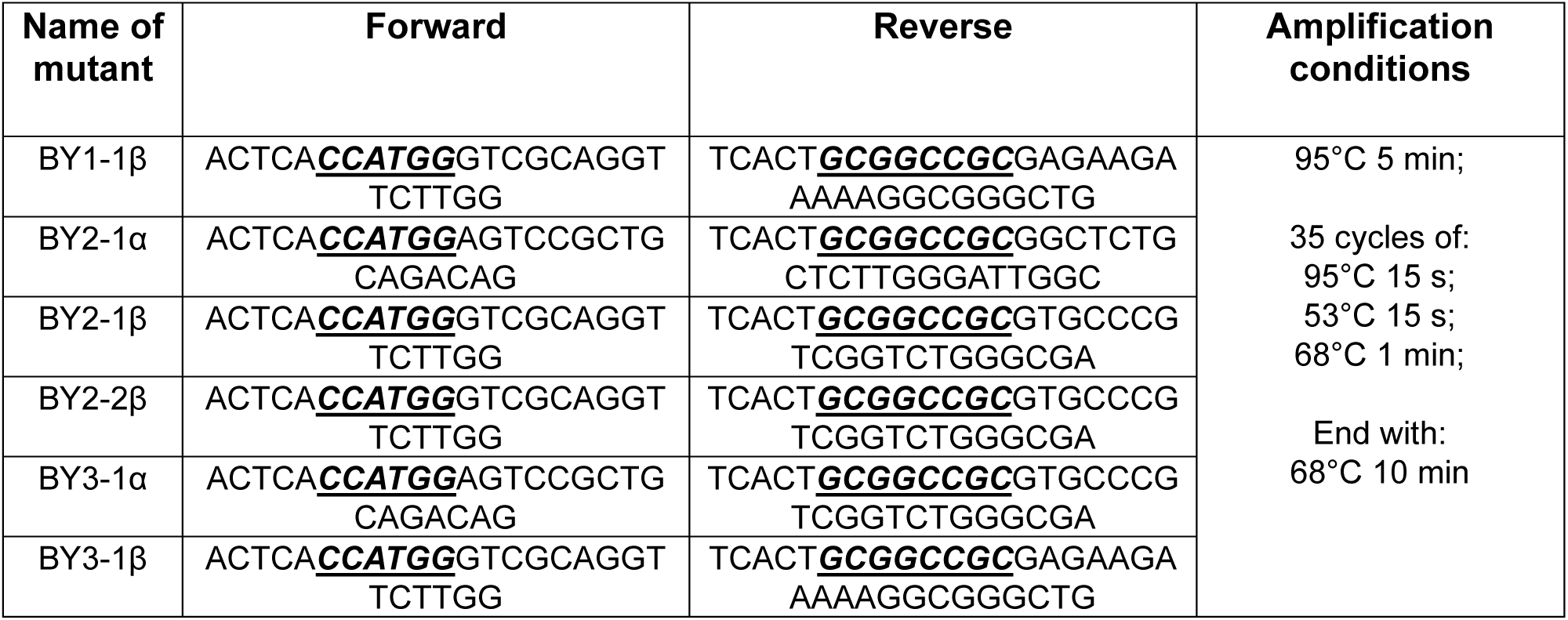
PCR primers used to amplify the cDNAs indicated in Figure 9.

The Tie-2-EGFP vector has been previously described ^52^ and was constructed by replacing the CMV early promoter in the pEGFP-N3 vector (Clontech/TaKaRa, Inc., San Jose, CA) with a 1.91 kb endothelial cell-specific Tie2 promoter fragment. Prior to transfection into tumor cells the vector was linearized with ApaL1. Stably transfected cells were selected in 400 μg/ml Geneticin (G418, ThermoFisher), pooled and used immediately for spheroid formation.

### Generation of primary HBs

HBs were induced in 6-8 wk old FVB/N mice (Jackson Labs, Bar Harbor, ME) as described previously using equal numbers of each sex. ^20, 21^ Briefly, 10 μg of each Sleeping Beauty (SB) vector encoding the stated combinations of B^del90^ Y^S127A^ and N^L30P^ (hereafter B,Y and N, respectively) were mixed at a 10:1 ratio with the pCMV-SB Transpose vector in a volume of 2 ml of sterile normal saline and then administered via hydrodynamic tail vein injection (HDTVI) over 5-10 sec. ^19–21^ To inactivate *Cdkn2a* in tumors, identical inocula contained 2 μg of Crispr/Cas9 pDG458 vectors (Addgene, Inc., Watertown, MA), each of which encoded gRNAs against 2 regions of the *Cdkn2a* exon 2 gene locus (Table 2 and Figure 1). Optimal gRNA sequences were identified using the CRISPOR selection tool. ^53^ Top strand gRNA sequences for pDG458-Vector (C1C2) were: C1: 5’-CGGTGCAGATTCGAACTGCGAGG-3’ and C2: 5’-GTCGTGCACCGGGCGGGAGAAGG-3’. Top strand gRNA sequences for pDG458-Vector (C3C4) were: C3: 5’-CTTGGGCCAAGTCGAGCGGCAGG-3’ and C4: 5’-TGCGATATTTGCGTTCCGCTGGG-3’. Top strand gRNA sequences for pDG458-Vector (C5C6), which were designed to specifically target exon 1α and inactivate p16^INK4A^ only were C5: 5’-AGGGCCGTGTGCATGACGTGCGG-3’ and C6: 5’ CTCCTTGCCTACCTGAATCGGGG-3’. The correct sequences and orientations of all gRNA-encoding pDG458 vectors were confirmed by dideoxy DNA sequencing.

**Figure 1.**
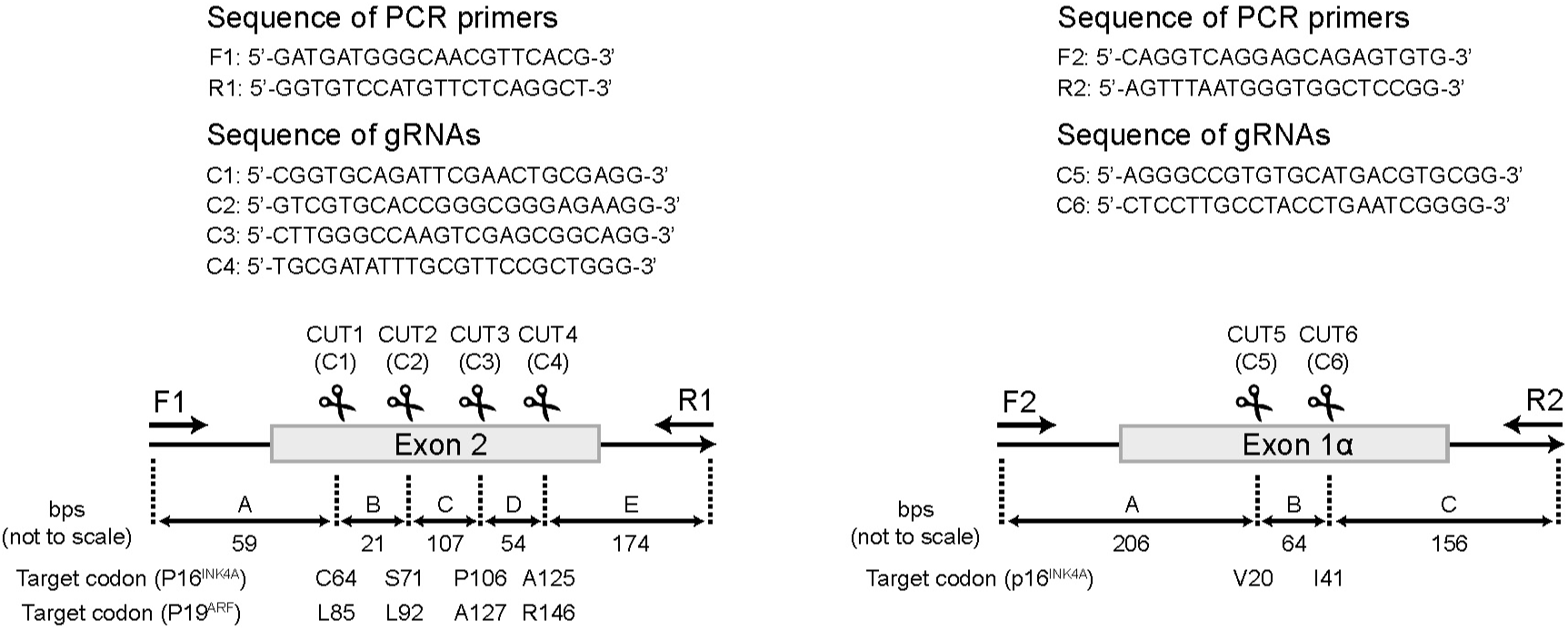
Sites of cutting for pDG458-Crispr/Cas9 vectors targeting the murine *Cdkn2a* locus. gRNAs C1C2 and C3C4 were specific for exon 2, which encodes both p16^INK4A^ and p19^ARF^ in alternate reading frames. ^40^ gRNAs C5C6 were specific for exon 1α, which encodes p16^INK4A^ only. Numbers shown beneath each cartoon indicate the distances (in bps) between each of the targeted sites and the PCR primers that were used to amplify both WT and mutant alleles (not to scale). High throughput sequencing of PCR products obtained using F1+R1 and F2+R2 primers (415 and 426 bps, respectively) was used to identify the precise Crispr/Cas9-induced mutations residing within various tumor cell populations (Supplementary File 1).

**Table 2.** Origin of immortalized HB cell lines.

| **Cell line name** | **Oncogenic drivers** | **pDG458**<br>**Crispr/Cas9**<br>**vector(s) used** | **Targeted Exon** |
| --- | --- | --- | --- |
| BY1 | B+Y | C1C2+C3C4 | 2 |
| BY2 | B+Y | C1C2+C3C4 | 2 |
| BY3 | B+Y | C1C2 | 2 |
| BY4 | B+Y | C1C2 | 2 |
| BY5 | B+Y | C1C2 | 2 |
| BY21 | B+Y | C5C6 | 1α |
| BY22 | B+Y | C5C6 | 1α |
| BY23 | B+Y | C5C6 | 1α |
| BYN1 | B+Y+N | C1C2+C3C4 | 2 |
| BYN2 | B+Y+N | C1C2+C3C4 | 2 |
| BYN3 | B+Y+N | C1C2+C3C4 | 2 |

### Immortalized HB cell lines

Upon reaching 8-10 g in size, tumors were removed from euthanized mice under sterile conditions, minced into multiple small fragments and washed several times in PBS. They were then digested with 0.1% trypsin for 30 min at 37C (Sigma-Aldrich, Inc. St Louis, MO) and further disrupted by gentle vortexing and vigorous pipetting followed by culturing in tissue culture plates containing standard D-MEM-10% FBS. Over the next several days tumor cells gradually attached to the surface of the plates but did not replicate. At this early stage of *in vitro* establishment, the cells were designated as “passage 0” (P0). After several additional weeks, multiple small colonies of densely packed replicating cells could be detected on each plate and were gradually expanded further. After 3-4 additional passages, aliquots were prepared for storage in liquid nitrogen and then used for subsequent studies.

### Tumorigenicity of immortalized HB cell lines

Monolayer cultures of BY or BYN tumor cells were trypsinized, washed twice in PBS and re-suspended in 0.2 of D-MEM (total of 10^6^ cells per inoculum unless otherwise stated). In some cases, Matrigel was added according to the directions of the supplier (Corning, Inc. Corning, NY). Cells were injected subcutaneously into the flanks of 4-6 wk old FVB/N mice or were administered by slow tail vein injection. In the former case, mice were monitored for the appearance of palpable tumors, which typically became detectable within 2-3 wks of injection. Tail vein-injected mice were sacrificed after 3 wks or earlier if noted to be in any distress. Regions of lung showing visible evidence of metastatic tumor growth were immediately fixed as described below.

### Tumor histology and immunohistochemistry

At the time of sacrifice, all tumors and relevant tissues were fixed in 4% paraformaldehyde, dehydrated in graded ethanol concentrations and then processed for H&E staining at the UPMC Children’s Hospital of Pittsburgh Core Histology Facility using standard procedures. For the detection of EGFP in tumor sections, small fragments of the subcutaneous tumors stably transfected with the Tie2-EGFP vector described above were embedded in O.C.T. compound (Thermo Fisher) After DAPI staining, fluorescence microscopy was performed with an Olympus Fluoview 1000 confocal microscope (Tokyo, Japan).

### SDS-PAGE, immunoblotting and immuno-staining

Livers, tumor tissues and cell lines were harvested as previously described and either lysed immediately for SDS-PAGE or snap-frozen in liquid nitrogen and stored at −80C until being further processed and analyzed. ^20, 21^ Standard lysis buffers were prepared in the presence of protease and phosphatase inhibitors according to the directions of the supplier (ThermoFisher). All immunoblotting was performed using PVDF membranes and semi-dry transfer conditions as previously described. ^20, 21^ All antibodies used for these studies at shown in Table 3.

**Table 3.** Antibodies Used in the Current Study.

| **Name of Antibody** | **Vendor** | **Catalog no.** | **Dilution used** |
| --- | --- | --- | --- |
| Anti-p16<sup>INK4A</sup> | Abcam | Ab211542 | 1:2000 |
| Anti-p19<sup>ARF</sup> | CST | 77184 | 1:500 |
| Anti-p19<sup>ARF</sup>(5-C3-1) | NovusBio | NB200-174 | 1:1000 |
| Anti-GAPDH | Sigma | G8795 | 1:10,000 |
| Anti-V5 | ThermoFisher | 46-07-05 | 1:5000 |
| Anti mouse HRP | CST | 7076 | 1:10,000 |
| Anti rabbit HRP | CST | 7074 | 1:5000 |

### Transcriptomic analyses

Raw RNA-seq data for human HBs were downloaded from the GEO database, specifically datasets GSE104766 ^10^ and GSE132219. ^8^ For mouse HB models, RNA-seq data were obtained from GEO datasets GSE130178 ^21^ and GSE157623. ^20^ All RNA-seq raw data were analyzed using nf-core/rnaseq version 3.12.0. ^54^ TPM values for genes or transcripts were used for quantification and heatmap generation. Heatmaps were created using the ComplexHeatmap R package. The transcript levels of human p16^INK4^ and p14^ARF^ from the TCGA cancer dataset were downloaded from UCSC Xena. ^55^ Correlations between p16^INK4^ and p14^ARF^ expression were visualized using scatter plots generated with the ggplot2 R package. Pearson’s correlation test was used to determine the strengths of the linear relationships. Gene set enrichment analysis (GSEA) was performed using the clusterProfiler R package.

For amplicon sequencing, DNAs purified from P0 and immortalized cell lines were amplified by PCR using different barcode-tagged primers. Equal amounts of purified PCR products were mixed and sequenced using the Illumina MiSeq platform, configured for paired-end 2 × 250 bp sequencing (Azenta Life Sciences, Inc., Chelmsford, MA). For analysis, the sequencing reads were first imported into CLC Genomics Workbench 24 (Qiagen), merged, and then de-multiplexed into groups based on the corresponding barcodes on the primers. The reads from each group were aligned to the mouse *Cdkn2a* locus with or without deletion (Supplementary File 1).

### Statistical analyses

R software v4.2.0 (R Foundation for Statistical Computing, Vienna, Austria) and GraphPad Prism v9.00 (Dotmatics, Inc. Boston, MA) were used for all statistical analyses. The ComplexHeatmap package was utilized for heatmap visualizations and the survminer package was used for curve plotting. The number of samples per group (n) for each experiment is indicated either in the figure legend or within the figure itself. Two-tailed, unpaired t-tests were used to evaluate significance between normally distributed populations, and a 2-tailed Mann-Whitney exact test was used for non-normally distributed populations.

## RESULTS

### Elevated expression of p16^INK4A^ and p19^ARF^ by murine and human HBs

An initial survey of experimental BY and BYN murine HBs showed that they expressed elevated levels of p16^INK4A^ and p19^ARF^ proteins relative to those in livers (Figure 2 A)^20^. Neither these nor BY tumors driven by 8 other patient-derived oncogenic B mutants showed evidence of p16^INK4A^ or p19^ARF^ transcript mutations (not shown). ^20, 21^ An analysis of 2 independent RNAseq studies comprising a total of 59 human HBs also showed significantly elevated mean levels of p16^INK4A^ and p14^ARF^ transcripts, all of which also encoded WT proteins (Figure 2 B&C). ^8, 10^ The more variable expression of these transcripts relative to those in murine HBs may have reflected the greater molecular heterogeneity of the human tumors as well as their more variable genetic backgrounds. Highly significant correlations were seen between p16^INK4A^ and p14^ARF^/p19^ARF^ transcripts in these HBs and in every set of human tumor samples from the TCGA database (Figure 3). Thus, experimental murine HBs, irrespective of their oncogenic driver background, appear to mimic the subset of human HBs with elevated levels of p16^INK4A^ and p14 ^ARF^/p19^ARF^.

**Figure 2.**
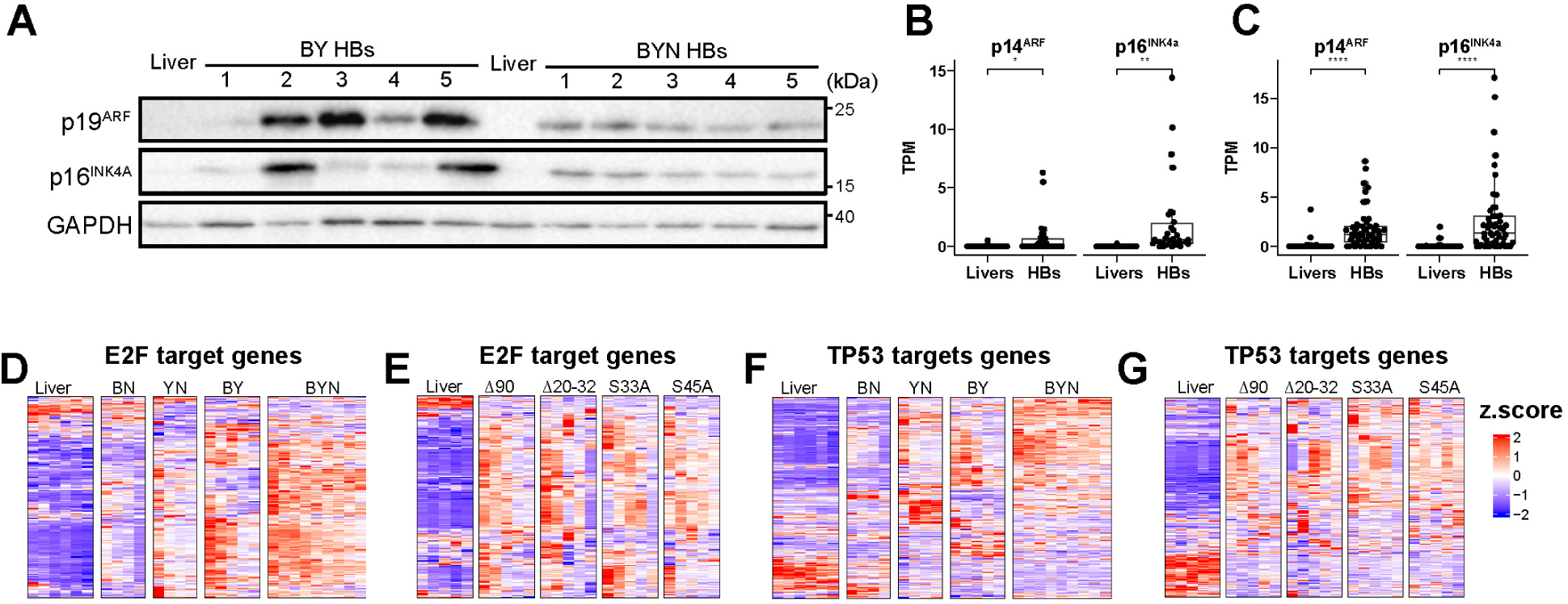
Increased expression of the *Cdkn2a* locus and its downstream target genes in murine and human HBs. (A). p16^INK4A^ and p19^ARF^ protein expression in control livers and HBs from murine BY and BYN HBs. (B). p16^INK4A^ and p14^ARF^ transcript levels in 25 human HBs. ^10^ (C). p16^INK4A^ and p14^ARF^ transcript levels in 34 human HBs. ^8^ (D). Heat maps of E2F target genes in age-matched control murine livers and the indicated HB types (E). Heat maps of E2F target genes in control murine livers and BY HBs generated by the indicated patient-derived B mutants as previously described.^21^ (F). Heat maps of TP53 target genes in age-matched control murine livers and the indicated HB types. (G). Heat maps of TP53 target genes in age-matched control murine livers and BY HBs generated by the indicated, patient-derived B mutants as previously described^21^.

**Figure 3.**
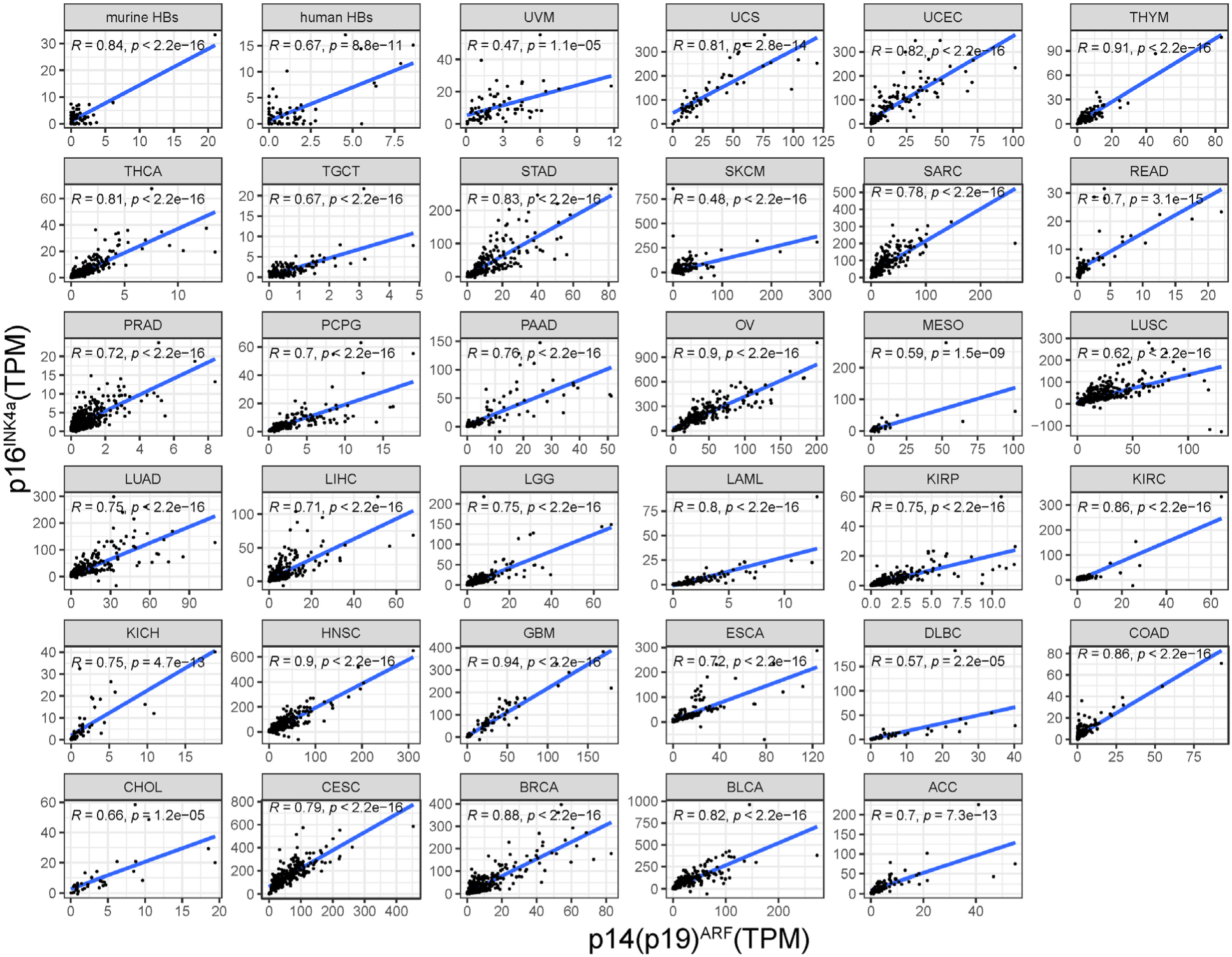
Positive correlations between p16^INK4A^ and p14^ARF^ transcripts across HBs and other human cancers. RNAseq data for murine and human HBs (first 2 panels: upper left) were compiled from Figure 2 B&C^8, 10^. RNAseq data for other, mostly adult, cancers were obtained from the UCSC Xena TCGA Hub. TPM was used as their absolute expression levels. Correlation coefficients and P values are shown in the upper left corner of each panel.

As a result of binding to Cdk4 and Cdk6 and inhibiting their intrinsic kinase activities, p16^INK4A^ indirectly maintains Rb in its active hypophosphorylated form. This in turn inhibits the G1-promoting activities of the E2F TF family thereby inducing cell cycle arrest and senescence. ^56, 57^ In parallel, p19^ARF^ blocks Mdm2’s E3 ubiquitin (Ub) ligase activity and TP53’s Ub-mediated degradation to further promote cell cycle arrest and apoptosis. ^39, 40, 58^ While the p16^INK4A^-Rb and p19^ARF^-TP53 pathways are distinct, they also cross-talk, with subsets of E2F transcriptional target genes contributing to apoptosis and subsets of TP53 target genes contributing to senescence. ^38, 59–62^ Despite the up-regulation of both p16^INK4A^ and p19^ARF^ in BY and BYN HBs (Figure 2 A), both their positive and negative target genes were regulated in directions that were the opposite of those predicted; moreover, similar, albeit less dramatic results were seen in BN and YN HBs and in BY HBs generated by several other B mutants (Figure 2 D-G, Figures 4 A & B and 5 A & B). ^21^ This seemingly aberrant form of E2F and TP53 target gene deregulation was also observed in human HBs where it was largely confined to the C2A subset of tumors with unfavorable outcomes (Figure 6 A&B and Figure 7 A&B). ^7, 8, 10^ Collectively, these studies suggested 2 possibilities. The first is that, despite strong activation of the *Cdkn2a* locus, its downstream transcriptional signaling and its promotion of senescence, apoptosis and cell cycle inhibition are overwhelmed by the strong oncogenic signaling of the B,Y and N drivers. Another less likely possibility is that both the Rb and TP53 pathways are inactivated and thus unresponsive to upstream signaling from p16^INK4^ or p19^ARF^ (*vide infra)*.

**Figure 4.**
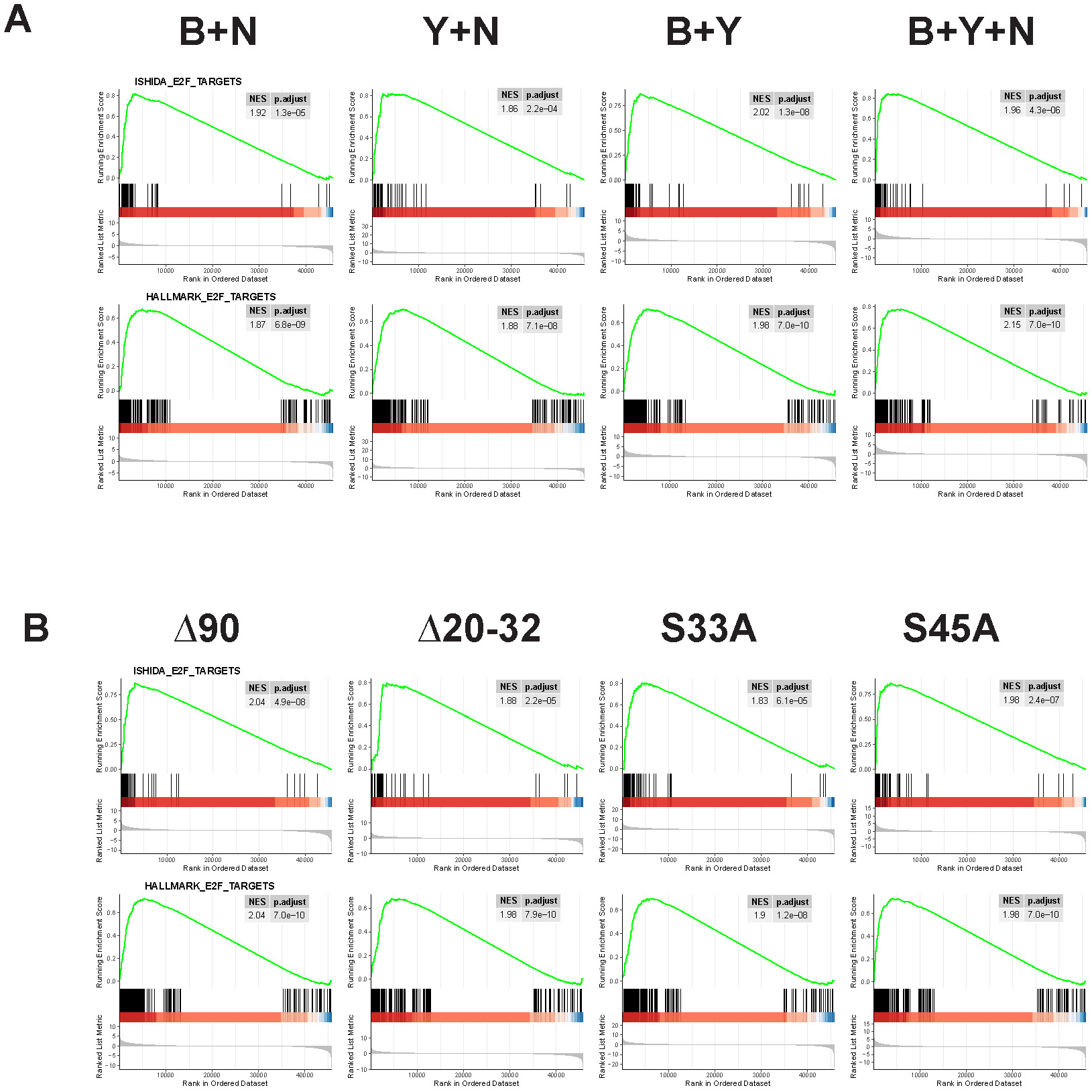
Individual GSEA plots for the E2F target gene groups shown in Figure 2 D and E. (A). Individual E2F target GSEA plots in BY and BYN tumors. (B). Individual E2F target GSEA plots in BY tumors with the indicated B mutations

**Figure 5.**
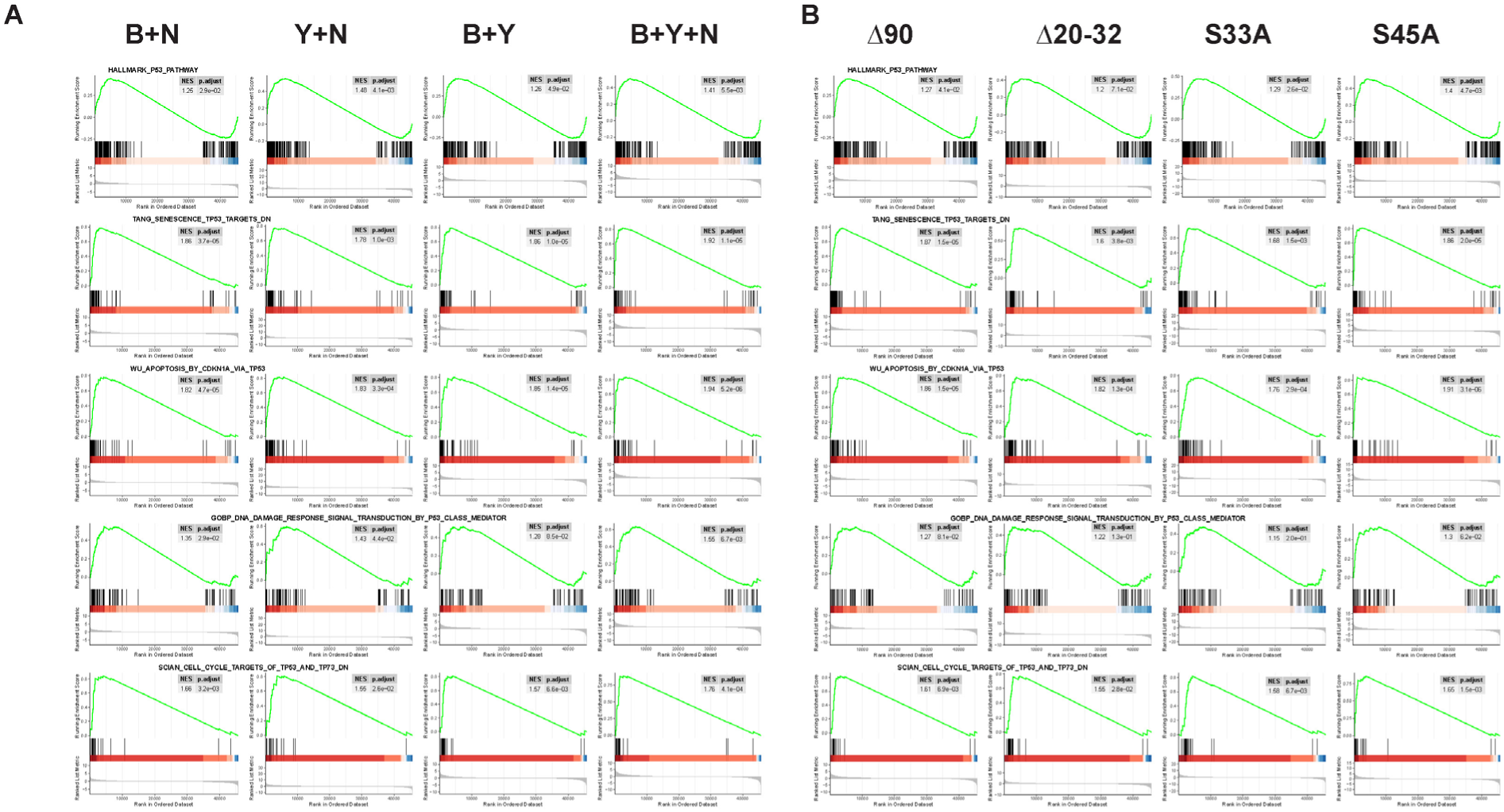
Individual GSEAs for the TP53 target gene groups shown in Figure 2 F and G. (A). TP53 target GSEA plots in HBs of the indicated sub-types. (B). TP53 target GSEA plots in BY tumors with different B mutations

**Figure 6.**
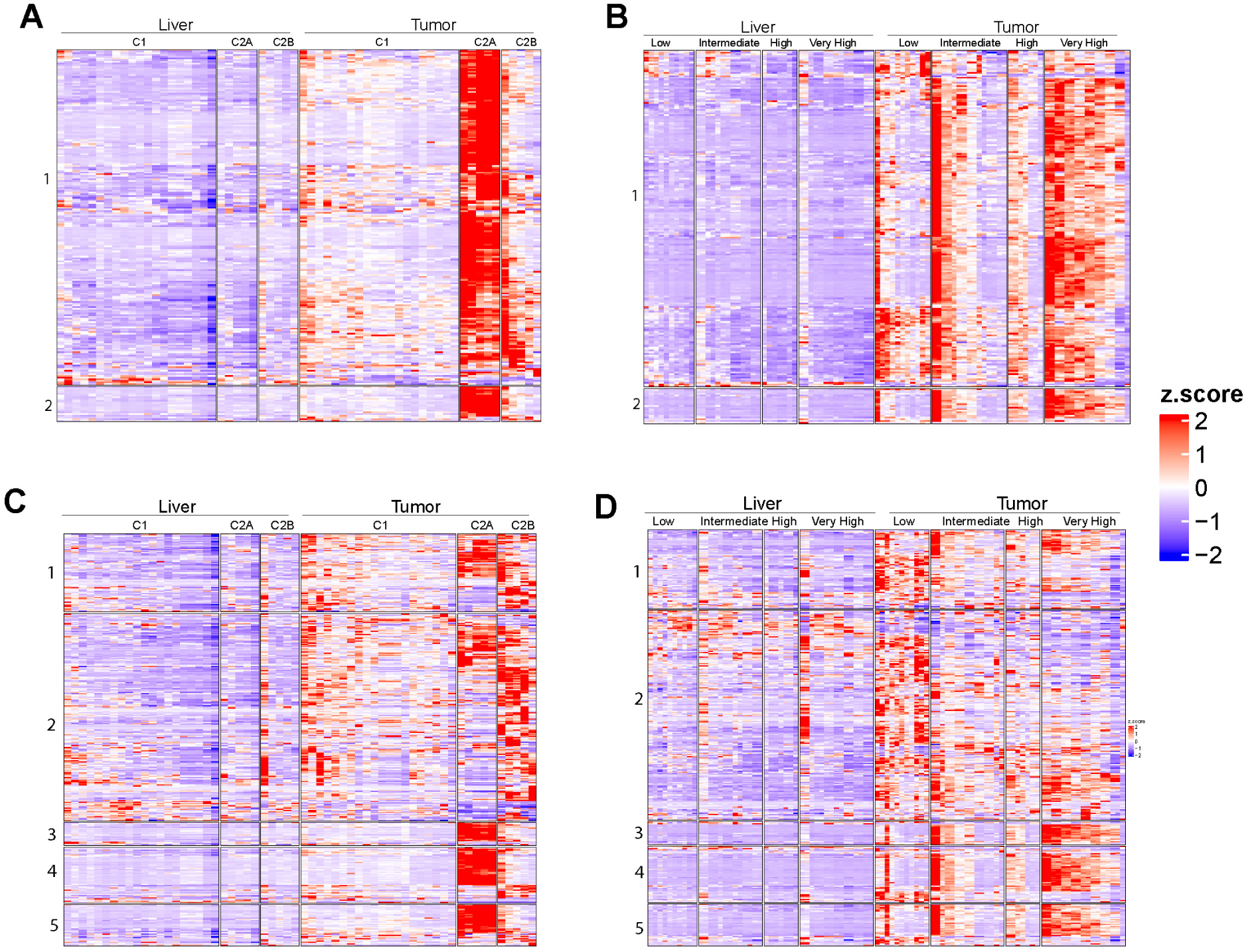
Dysregulation of E2F and TP53 target genes. (A). Heatmaps of E2F target gene expression relative to that of matched normal liver samples in 25 human HBs from GSE104766 ^10^. Gene sets 1 and 2 are from the MSigDB gene sets HALLMARK_E2F_TARGETS and ISHIDA_E2F_TARGETS, respectively. (B). Heatmaps of E2F target gene expression relative to that of matched normal liver samples in 33 human HBs from GSE132219 ^8^. The same gene sets as shown in A are used. (C). Heatmaps of TP53 target gene transcript levels relative to those of matched normal liver samples in the samples shown in A. TP53 target gene sets are derived from the following MSigDB gene sets: **1** GOBP_DNA_DAMAGE_RESPONSE_SIGNAL_TRANSDUCTION_BY_P53_CLASS_MEDIATOR, **2** HALLMARK_P53_PATHWAY, **3** SCIAN_CELL_CYCLE_TARGETS_OF_TP53_AND_TP73_DN, **4** TANG_SENESCENCE_TP53_TARGETS_DN, and **5** WU_APOPTOSIS_BY_CDKN1A_VIA_TP53. (D). Heatmaps of TP53 target genes relative to those of matched normal liver samples in the samples shown in B. The same gene sets as shown in C are used.

**Figure 7.**
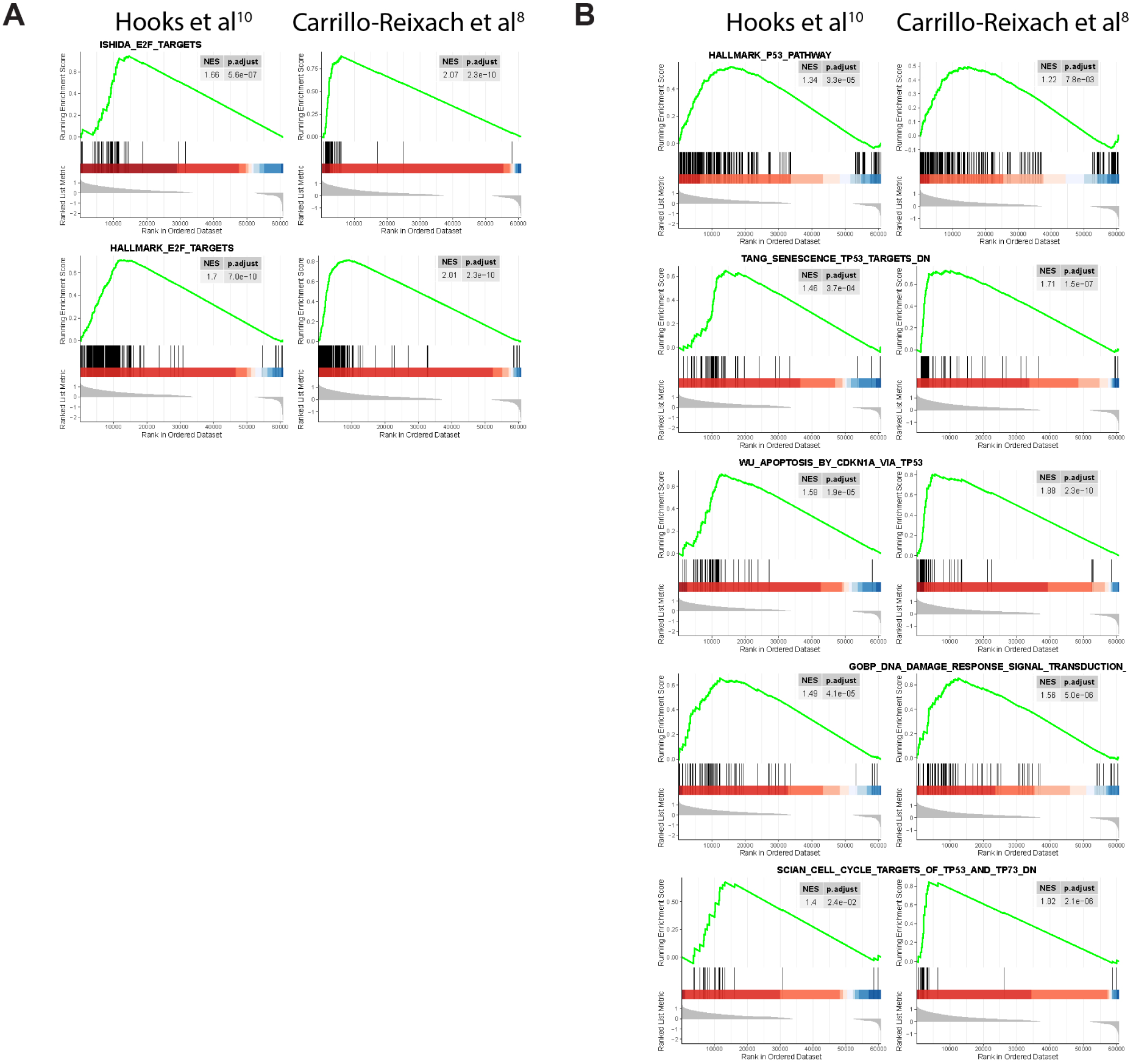
Individual GSEA plots for the genes depicted in Figure 6 from the C2A subset of human HBs. (A). Individual E2F target GSEA plots for the transcripts depicted in Figure 6 A and B. (B). Individual TP53 target GSEA plots for the transcripts depicted in Figure 6C and D.

### Efficient establishment of immortalized murine HB cell lines after *in vivo Cdkn2a* locus inactivation

Our previous attempts to establish immortalized cell lines from murine HBs induced by different B, Y and N combinations proved unsuccessful on over 30 occasions (H. Wang and E. Prochownik, unpublished). However, the above findings suggested that concurrent inactivation of the *Cdkn2a* locus during the formative stages of tumorigenesis might allow for more rapid *in vivo* tumor growth and *in vitro* establishment by eliminating senescence and apoptotic barriers that potentially restrain proliferation and immortalization. ^38–40, 49, 63, 64^ To test this, we generated BY and BYN HBs in FVB/N mice utilizing various combinations of 3 pDG458-Crispr/Cas9 vectors, each of which encoded 2 different gRNAs directed against the *Cdkn2a* locus (Table 2). The C1C2 and C3C4 vectors were each designed to target 2 different regions of exon 2’s common p16^INK4A^ and p19^ARF^ coding region and the C5C6 vector was designed to target 2 regions of the p16^INK4A^-specific 1α exon (Figure 1). The 30 BY HBs generated with these vectors were sufficient to conclude that each one, whether delivered alone or together, accelerated BY tumor growth rates and shortened median survival (Figure 8 A). Because only 3 BYN+C1C2+C3C4 tumors were generated and because control BYN tumors already grew at an exceedingly rapid rate ^20^, it was not possible to ascertain whether targeting *Cdkn2a* significantly accelerated their growth. Examination of a randomly chosen subset of primary tumors reveled that some expressed either no or extremely low levels of p16^INK4A^ or p19^ARF^, with the latter likely originating from non-hepatocytes, from tumor cells in which the *Cdkn2a* locus remained intact or from hepatocytes with p16^INK4A^ and p19^ARF^ mutations comprising small indels or point mutations that could not be resolved from the WT proteins (Figure 8B). Several of the HBs also expressed p16^INK4A^ and p19^ARF^ proteins that migrated more rapidly than WT proteins. Of the 3 tumors, with *Cdkn2a* exon 1α-specific targeted by the C5C6 gRNA combination, one expressed no p16^INK4A^, 2 expressed very low levels and one appeared to express no p19^ARF^.

**Figure 8.**
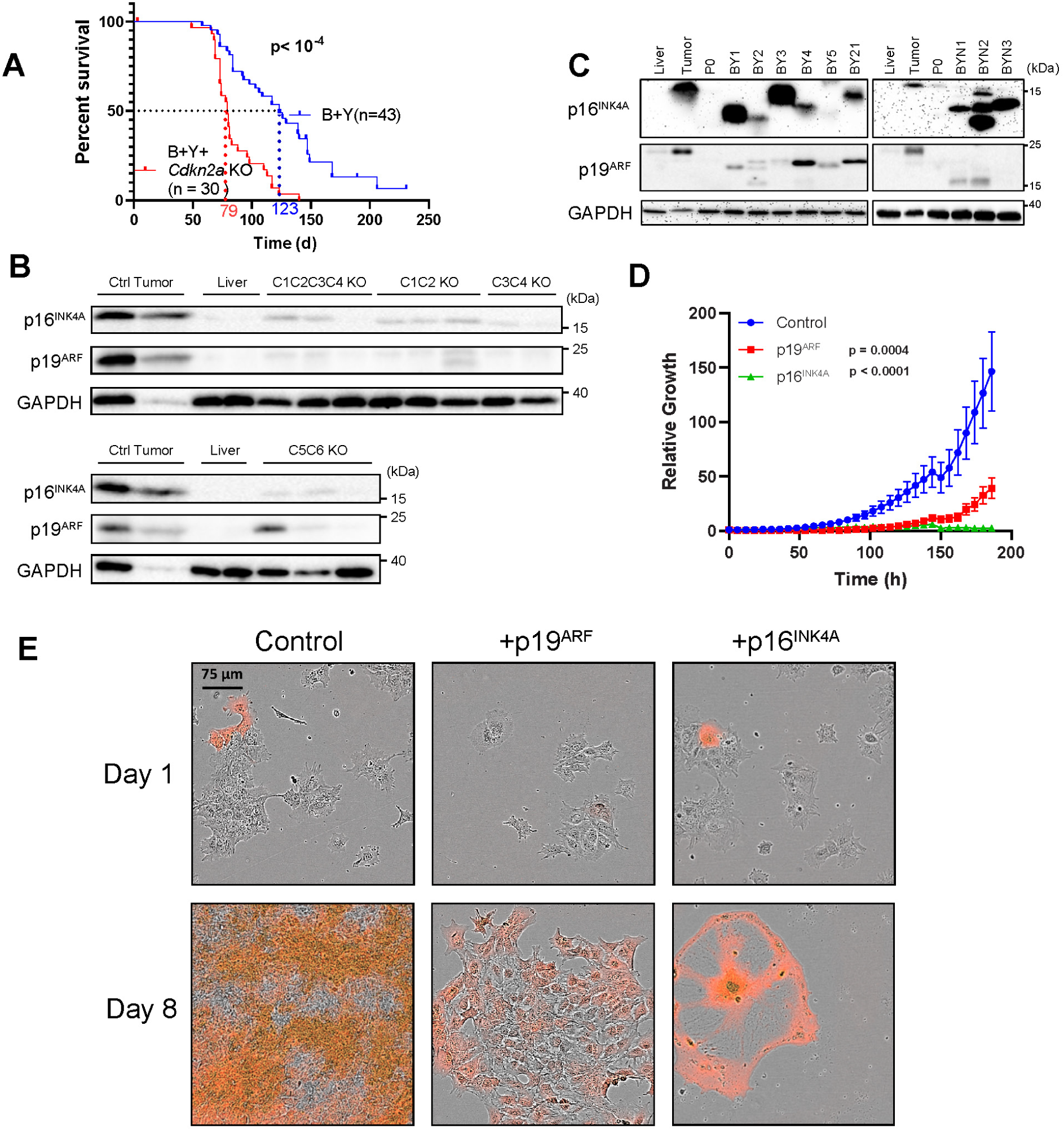
*In vivo* targeting of the *Cdkn2a* locus in murine HBs accelerates tumor growth and allows for the establishment of immortalized cell lines that can be inhibited by re-expressing p16^INK4A^ or p19^ARF^. (A). Kaplan-Meier survival curves of mice with BY HBs (control group) and those with BY HBs in which the *Cdkn2a* exon 2 locus was targeted with the pDG458 Crispr/Cas9 vectors listed in Table 2 and Figure 1. Seven-9 mice from each group were combined and are generically designated as “BY+*Cdkn2a* KO”. (B). Markedly reduced p16^INK4A^ and p19^ARF^ protein expression in randomly selected BY+*Cdkn2a* KO primary HBs from A. Protein levels are compared to those from control BY tumors and normal livers. (C). p16^INK4A^ and p19^ARF^ proteins in the indicated BY+*Cdkn2a* KO and BYN+*Cdkn2a* KO cell lines. “P0” lysates were from HB cells that had been recently plated *in vitro* but did not proliferate and had not yet generated immortalized clones. Liver (L) and control BY HB (T) lysates were included as controls. (D). Growth curves of BY1 cells following *in vitro* stable transfection with a control empty SB vector or with those encoding WT p16^INK4A^ or p19^ARF^. Each of the other cell lines showed similar growth kinetics (bot shown). (E). Typical microscopic appearance of the cells from D following stable transfection of the indicated SB vectors which also expressed dTomato. Note the presence of an extremely large and senescent-like p16^INK4A^ expressing cell.

In contrast to our previous failure to establish immortalized HB cell lines from tumors with intact *Cdkn2a* loci, we readily generated 8 BY and 3 BYN cell lines in 11 attempts from tumors in which *Cdkn2a* had been targeted, with each having been maintained continuously for at least 4 months. As was true of the Crispr/Cas9-targeted primary tumors (Figure 8B), we detected very little if any p16^INK4A^ or p19^ARF^ protein in these non-proliferating HB tumor cells following their initial *in vitro* plating (“P0” populations) Figure 8C). Following the outgrowth of immortalized and highly proliferative tumor cell populations from these cells, however, we identified what in most cases were truncated p16^INK4A^ and/or p19^ARF^ proteins (Figure 8C). Collectively, these studies indicate that immortalized cell lines can be readily and routinely generated from BY and BYN HBs when *Cdkn2a* is concurrently inactivated. Finally, the enforced expression of WT p16^INK4A^ or p19^ARF^ strongly inhibited the *in vitro* growth of these tumor cells (Figure 8D) and, in the former case, caused many of them to become dramatically enlarged and senescent-like in appearance (Figure_8E). These results show that achieving an immortalized state requires the disabling of the *Cdkn2a* locus. It also showed that pathways downstream of p16^INK4A^ and p19^ARF^ remained intact given that the re-expression of either TS was sufficient to achieve virtually total growth suppression.

### Both positive and negative selection of p16^INK4A^ and p19^ARF^ mutants are associated with *in vitro* outgrowth of HB cell lines

To determine the identities and frequencies of the *Cdkn2a* mutations in primary tumors, whether they were associated with clonal expansion or regression during subsequent immortalization and whether they were recurrent, we performed high-throughput DNA sequencing of PCR-amplified exon 2 or exon 1α products from 9 P0-immortalized cell line pairs. To ensure an unbiased representation from all regions of each HB, DNAs were isolated from P0 monolayer cultures several days after plating and well before immortalized tumor cell colony outgrowth was observed. The sequences of the documented *Cdkn2a* mutations were then compared to those of the immortalized cell lines that eventually emerged from and replaced the non-dividing P0 populations. After excluding large deletions, an average of 37 different *Cdkn2a* mutations (range 18-57) were identified among all P0 populations, which were reduced by ∼40% following immortalization (average = 22, range 11-33 P=0.006) (Table 4). This indicated that many of the *Cdkn2a* mutations originally generated in primary tumors conferred little-no survival benefit and thus were subject to negative selection. Although this analysis could not determine the total number of independently arising clones, it did set lower limits on them.

**Table 4.** Total number of unique *Cdkn2a* locus mutations identified in each P0 and immortalized cell population.

| **ID** | **Type** | **Total Number of Reads** | **Number of Unique Reads\*** | **Immortalized: P0 ratio** |
| --- | --- | --- | --- | --- |
| BY1 | P0 | 1874 | 39 | 0.46 |
| BY1 | Immortalized | 2819 | 18 |  |
| BY2 | P0 | 3433 | 29 | 0.86 |
| BY2 | Immortalized | 3291 | 25 |  |
| BY3 | P0 | 1126 | 33 | 0.55 |
| BY3 | Immortalized | 3943 | 18 |  |
| BY4 | P0 | 2190 | 27 | 0.70 |
| BY4 | Immortalized | 1374 | 19 |  |
| BY5 | P0 | 1006 | 35 | 0.40 |
| BY5 | Immortalized | 287 | 14 |  |
| BY21 | P0 | 113 | 18 | 0.61 |
| BY21 | Immortalized | 73 | 11 |  |
| BYN1 | P0 | 2978 | 48 | 0.63 |
| BYN1 | Immortalized | 12317 | 30 |  |
| BYN2 | P0 | 3558 | 57 | 0.58 |
| BYN2 | Immortalized | 12630 | 33 |  |
| BYN3 | P0 | 3722 | 41 | 0.54 |
| BYN3 | Immortalized | 4723 | 22 |  |

\* Does not include large deletions.

All 9 P0 and immortalized cell line pairs possessed a relatively small number of pre-eminent fusion and/or deletion mutations (Figure 9A and Supplementary File 1). Notable examples of this were observed in the BY1, BY2, and BY4 P0 cells. Their retention by the immortalized populations implicated them as either being neutral or conferring an equivalent survival and/or proliferative benefit to both populations. In other cases, minor mutations detected in P0 populations expanded significantly during subsequent immortalization, with particularly notable examples occurring in BY3 and BYN2 cells. Negative selection of certain mutants in immortalized cells was also observed, as for example in BY1, BY3 and BYN1 tumors.

**Figure 9.**
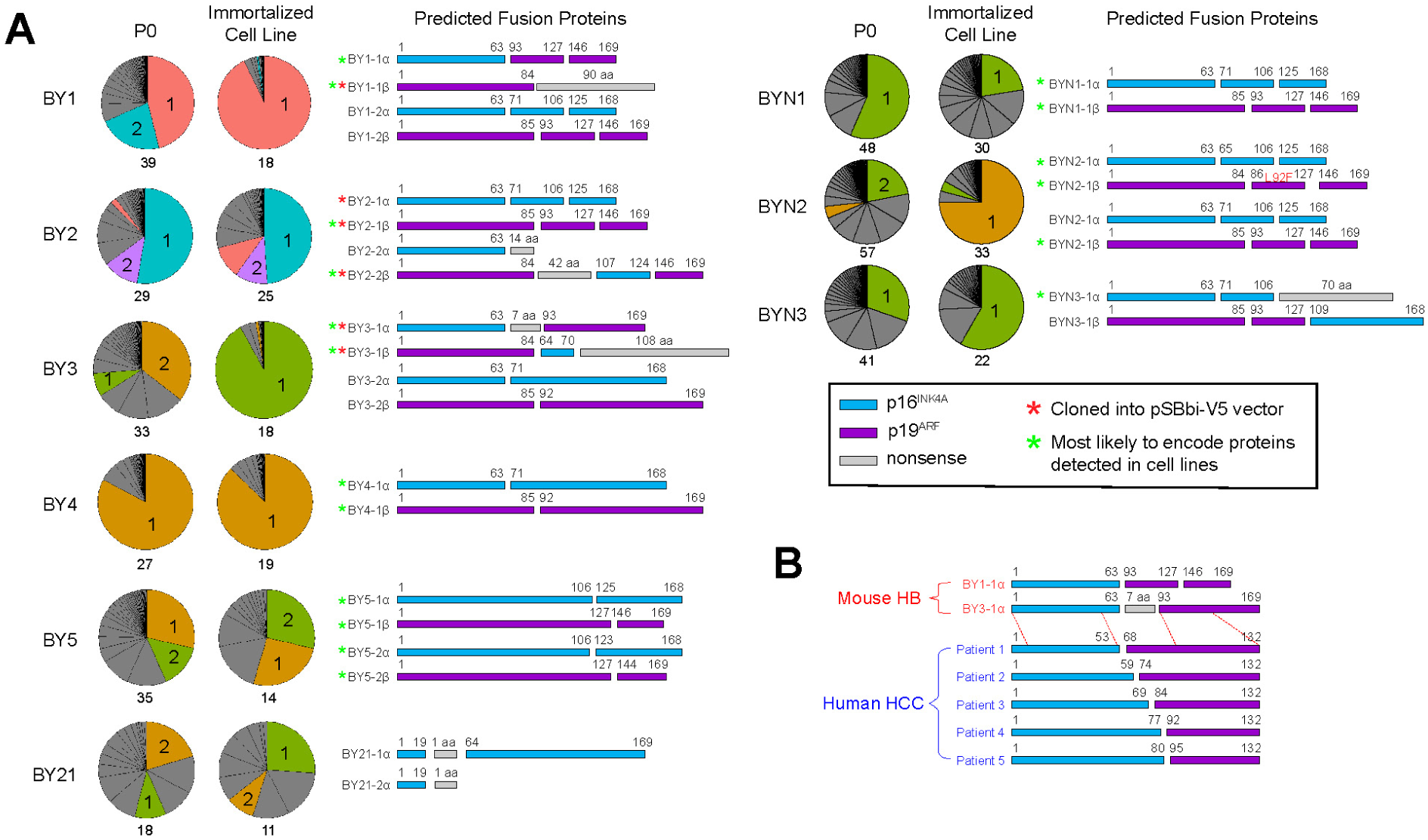
Mutational neutrality or selection of *Cdkn2a-*targeted loci in P0 and immortalized hepatocyte populations. (A). Pie charts indicating the frequency with which various *Cdkn2a* mutations were identified in P0 and immortalized cell populations after excluding deletions larger than C2-C3 (Figure 1). (See Supplementary File 1 for the actual sequences and their frequencies). Numbers below each pie chart indicate the total number of distinct mutations that could be identified. Colored segments indicate mutants that were of particular prominence and/or interest in one or both populations. Shown to the right of the pie charts are renditions of the proteins predicted to be encoded by the colored segments that would be translated from transcripts initiated from either *Cdkn2a* exon 1α or 1β. All transcripts depicted were expressed as they could be amplified by RT-PCR. Red asterisks indicate transcripts that were amplified and cloned into the pSBbi-RP-V5 SB expression vector for further evaluation. Green asterisks indicate the transcripts that were most likely responsible for encoding the proteins shown in Figure 8 C. (B). 5 HCCs from the COSMIC liver cancer database express p16^INK4A^-p14^ARF^ fusion transcripts that closely resemble some of those generated by the deliberate targeting of the *Cdkn2a.* Dotted lines indicate the locations of homologous residues in the human and murine sequences. The murine fusion proteins shown are taken from panel A.

A more detailed examination of the mutant transcripts that were predicted to be expressed indicated that they encoded a combination of p16^INK4A^ and p19^ARF^ in-frame deletion mutants and p16^INK4A^-p19^ARF^ fusions (Figure 9A). Examples of the former included those encoded by the BY1-2α and BY3-2β transcripts whereas examples of the latter included those encoded by BY1-1α, BY21-1α and BYN3-1α, with the latter 2 containing short (1-7 amino acids) nonsense insertions. Transcripts encoding truncated proteins with nonsense extensions were seen in the case of BY1-1β whereas complex mutations encoding various combinations of indels, fusions, truncations and extensions were observed in the cases of BY3-1β, BYN1-1α and BYN3-1β. Finally, several tumors and/or their progeny were found to have mutations that were either identical to or closely related to those generated in other HBs, such as those for BY1-2β/BY2-1β and BYN1-2α/BYN2-2α. Proteins of the sizes predicted by *in silico* translation of the sequences shown in Figure 9A matched those actually observed in the cell lines using antibodies directed against p16^INK4A^ and p19^ARF^ although more than one of these were seldom detected (Figure 8C). Collectively, these findings show that P0 and immortalized cells contain p16^INK4A^ and p19^ARF^ fusions or deletions that are selected for only in the primary tumor, only in the immortalized cells derived from these tumors or in both the tumors and the immortalized cells.

A survey of 3377 human liver cancers from the COSMIC cancer database identified 5 tumors (all HCCs) with mutations in the *CDKN2A* locus that generated p16^INK4A^-p14^ARF^ fusion transcripts with striking similarities to the BY1-1α and BY3-1α murine HB fusions (Figure 9B). These human transcripts were predicted to encode proteins in which the first 53-80 amino acids of p16^INK4A^ were fused in-frame to residues 68-95 of p14^ARF^. The fact that these closely related transcripts were identified in both adult- and pediatric-type liver cancers from 2 different species and that the BY1-1α and BY3-1α transcripts were markedly enriched during the transition to immortalization suggested either that they play an active role in the outgrowth of these cell populations or that they are less repressive than other mutations that were lost over the course of time.

Finally, we identified 158 additional tumors from the above database with *CDKN2A* mutations predicted to encode p16^INK4A^-p14^ARF^ fusion proteins similar to those identified in the above-mentioned liver cancers. The most common cancers associated with these included those of the esophagus (20 cases), skin (20 cases) and lung (14 cases). As with those depicted in Figure 9B, nearly all involved residues 53-80 of p16^INK4A^ and residues 68-95 of p14^ARF^. While the vast majority of these were unique, one recurrent mutation, identical to that detected in HCC Patient 4 was identified 31 times (Figure 9B) (https://cancer.sanger.ac.uk/cosmic/mutation/overview?id=146577689.

To determine whether p16^INK4A^-p19^ARF^ fusion proteins could differentially influence the growth of immortalized HB cell lines, we amplified the cDNAs encoding some of the more prominent mutant proteins shown in Figure 9A, cloned them into the pSBbi-RP-V5 SB vector and expressed them transiently in BY21 cells. Immunoblotting with an anti-V5 antibody confirmed that each vector encoded a protein of the predicted size while allowing the relative levels of each to be compared (Figure 10A). Subsequent stable expression of these mutants in BY21 cells caused variable degrees of growth suppression (Figure 10B).

**Figure 10.**
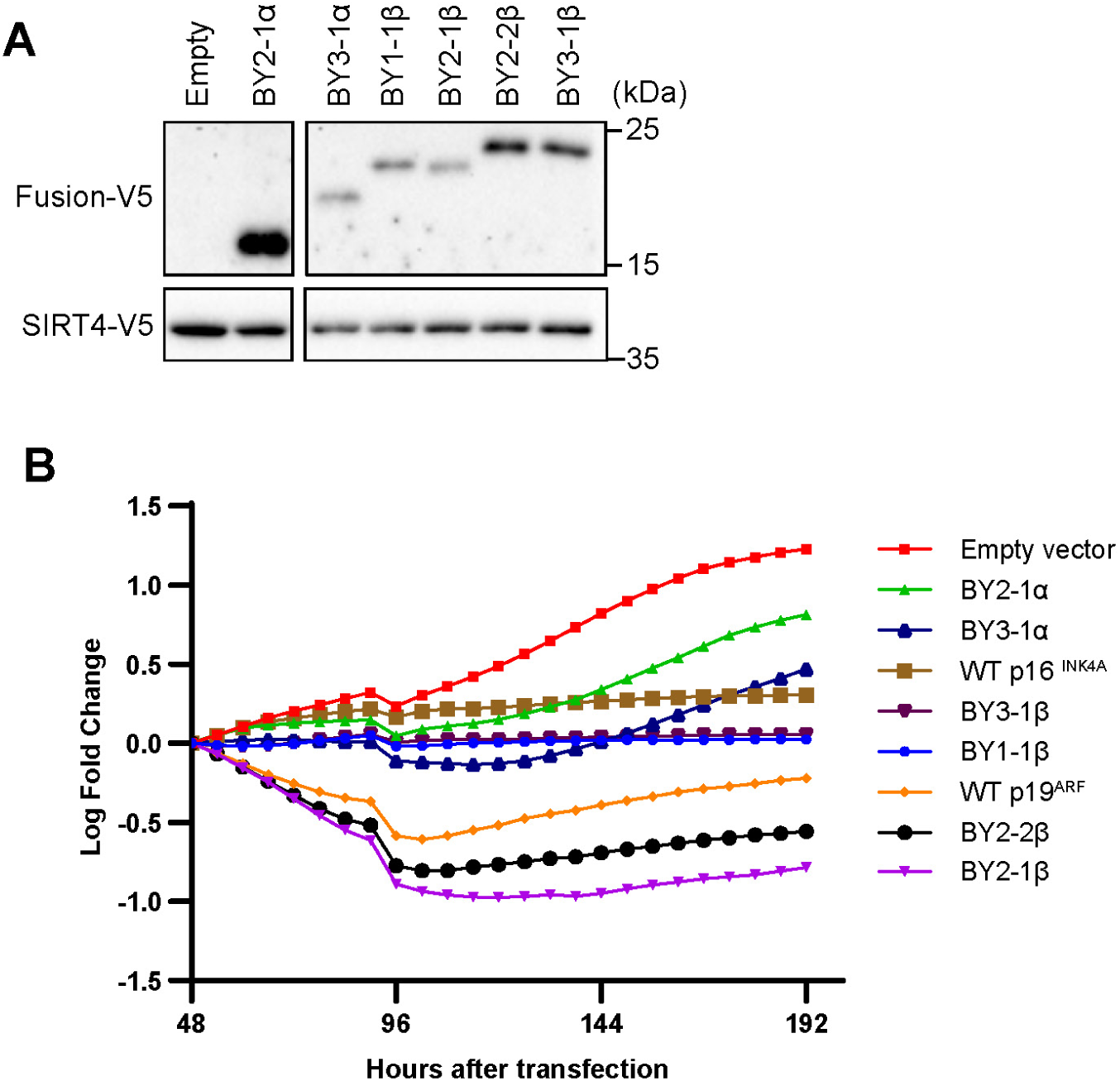
Mutant p16^INK4A^ and p19^ARF^ proteins impact HB cell behaviors. (A). The indicated V5 epitope-tagged fusion proteins from Figure 5A (asterisks) were transiently expressed in BY21 cells and detected by immunoblotting in whole cell lysates using an anti-V5 tag antibody. To control for any differences in transfection efficiencies, each plasmid was co-transfected with an equal amount of a vector that encoded a V5-epitope-tagged Sirt4 protein. (B). The proteins shown in A were stably expressed in BY1 cells as indicated and maintained in puromycin-containing medium to allow for selection of dTomato+ cells. Control cells were similarly transfected with the empty pSBbi-RP-V5 SB vector. Cell counts were then performed as described for Figure 4D.

During the course of the above studies, we were successful in generating one immortalized cell line originating from a single clone derived from a BY tumor that had not been targeted with a *Cdkn2a* Crispr vector. Sequence analysis of the clone’s *Cdkn2a* coding exons showed them to be wild-type despite the fact that neither p16^INK4A^ nor p19^ARF^ proteins could be detected (K. Chen and E. Prochownik, unpublished). We conclude that this clone is likely related to the subset of human HBs depicted in Figure 2B whose *Cdkn2a* loci are likely to be silenced as a result of promoter hypermethyation ^44^.

### Most immortalized BY and BYN cell lines retain their morphologic features and tumorigenic behaviors

Like the primary crowded fetal-like HBs from which they originated ^20^, all 11 BY and BYN cell lines appeared morphologically similar. Individual cells were mostly mononuclear, with prominent and often multiple nucleoli and low nuclear:cytoplasmic ratios (Figure 11A). ^20^ Despite the similarities among their individual cell populations, those from the 3 BYN groups were more inclined to grow in discrete colonies, which tended to form within 3-4 days of plating even from otherwise well-dispersed and lightly seeded single cell populations. 10 of the 11 cell lines formed rapidly growing subcutaneous tumors in the flanks of FVB/N mice even in the absence of a facilitating basement membrane extract such as Matrigel (Figure 11B). However, even the inclusion of Matrigel did not support tumor formation by the BY2 cell line. Tumors generated from a cell population containing equal numbers of EGFP-tagged BY2 cells and untagged BY1 expressed no EGFP, thus indicating that the former cells’ lack of tumorigenicity was likely cell intrinsic (not shown). In the remaining 10 cases, the subcutaneous tumors which did form retained the crowded fetal HB-like histology of the original tumors. ^19–21^ Following tail vein injection, at least 6 of these cell lines could also grow as metastasis-like deposits within the lung parenchyma (Figure 11C). Additionally, in a mouse in which a BY1-subcutaneous tumor was allowed to persist for 6-7 weeks rather than the usual 3-4 weeks, we also observed pulmonary metastases, indicating that at least some of our HB cell lines can spontaneously metastasize from distant sites (Figure 11D). The HB lines described here thus generally maintained the appearance and behavior of the primary tumors from which they originated while, in most cases, also providing models of metastatic disease.

**Figure 11.**
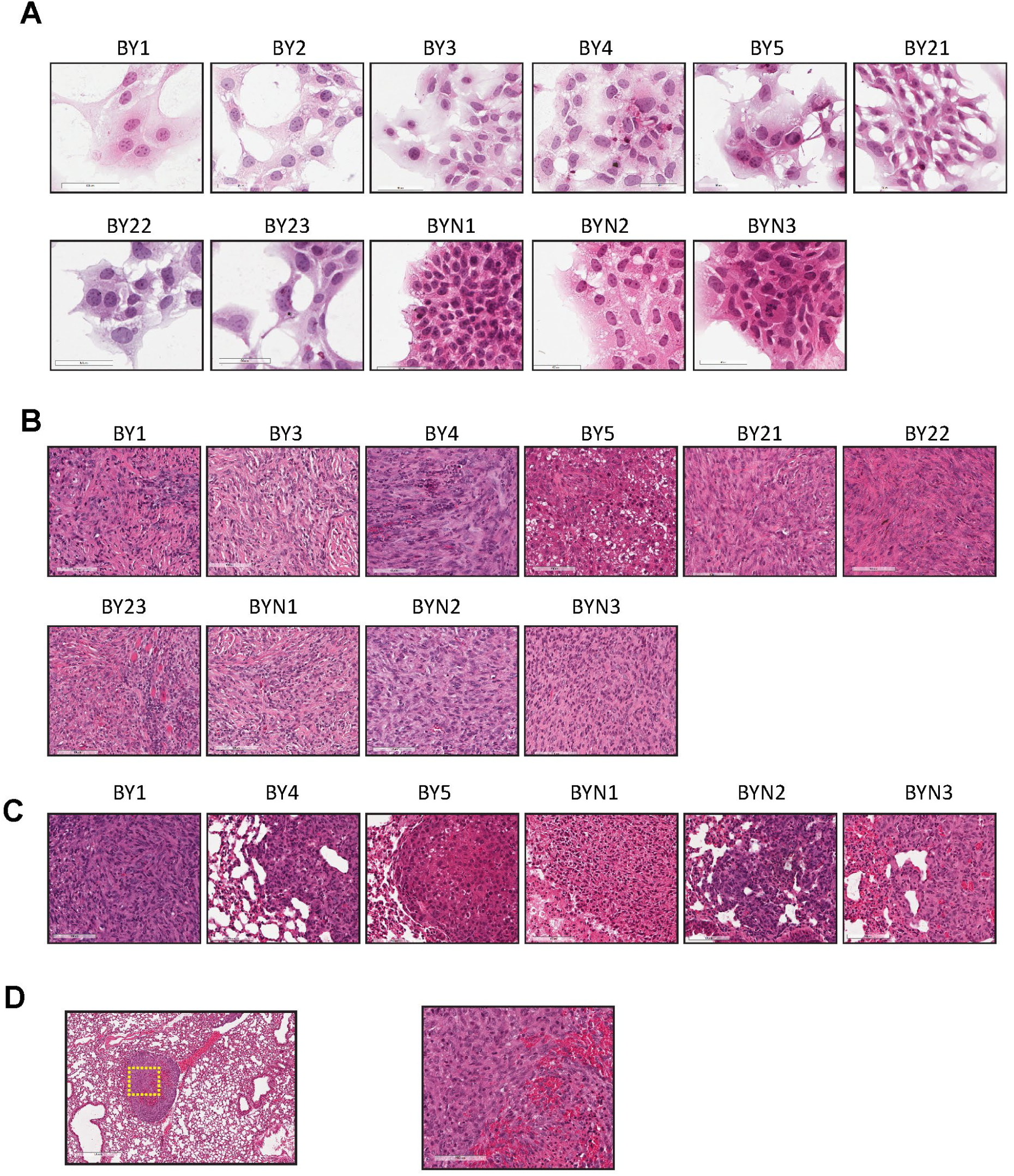
Histologic appearance of Immortalized cell lines and tumors. (A). H&E-stained monolayers of the indicated cell lines grown on glass coverslips. (B). H&E-stained paraffin-embedded sections of subcutaneous tumors generated by the indicated cell lines. (C). H&E-stained paraffin-embedded sections of tumors metastatic to lung following tail vein injection of the indicated cell lines. (D). Left: low-power image of a spontaneous pulmonary metastasis in a mouse subcutaneously inoculated with BY1 cells 7 weeks earlier. Right: higher power image of the region indicated by the box in the left-most panel.

### HB cell lines reversibly form dynamic spheroids and tumors whose hypoxic interiors are permissive for endothelial cell-specific gene expression

Tumor organoids and spheroids are 3-dimensional cellular aggregates that can be maintained *in vitro* where they display properties and behaviors more akin to those of actual tumors than do cells maintained as monolayers. ^6, 65–67^ Organoids, including those derived from human HBs, are more complex than spheroids in that they contain additional non-transformed cells such as fibroblasts, endothelial cells and immune cells that may be components of the primary tumor or are deliberately added *in vitro.* ^68, 69^ Tumor spheroids, in contrast, are derived only from tumor cells and are generally easier to maintain long-term. ^65, 70^ To determine whether HB cell lines could form spheroids, single cell suspensions were seeded into polypropylene test tubes or plastic bacterial-grade Petri dishes. In the former case, cells remained unattached, whereas in the latter case, they initially formed monolayers that appeared similar to those formed on standard tissue culture dishes (Figure 12A). Under at least one of these conditions, all 11 cell lines formed organoid-like aggregates within 3-7 days and could be maintained in this manner for >3 weeks (Figure 12 A and B). That the cells remained highly viable was demonstrated by re-plating the spheroids onto standard tissue culture plates where they rapidly reassumed monolayer growth (Figure 12A). Individual spheroids were also highly dynamic and could fuse to form single hybrid spheroids (Figure 12C).

**Figure 12.**
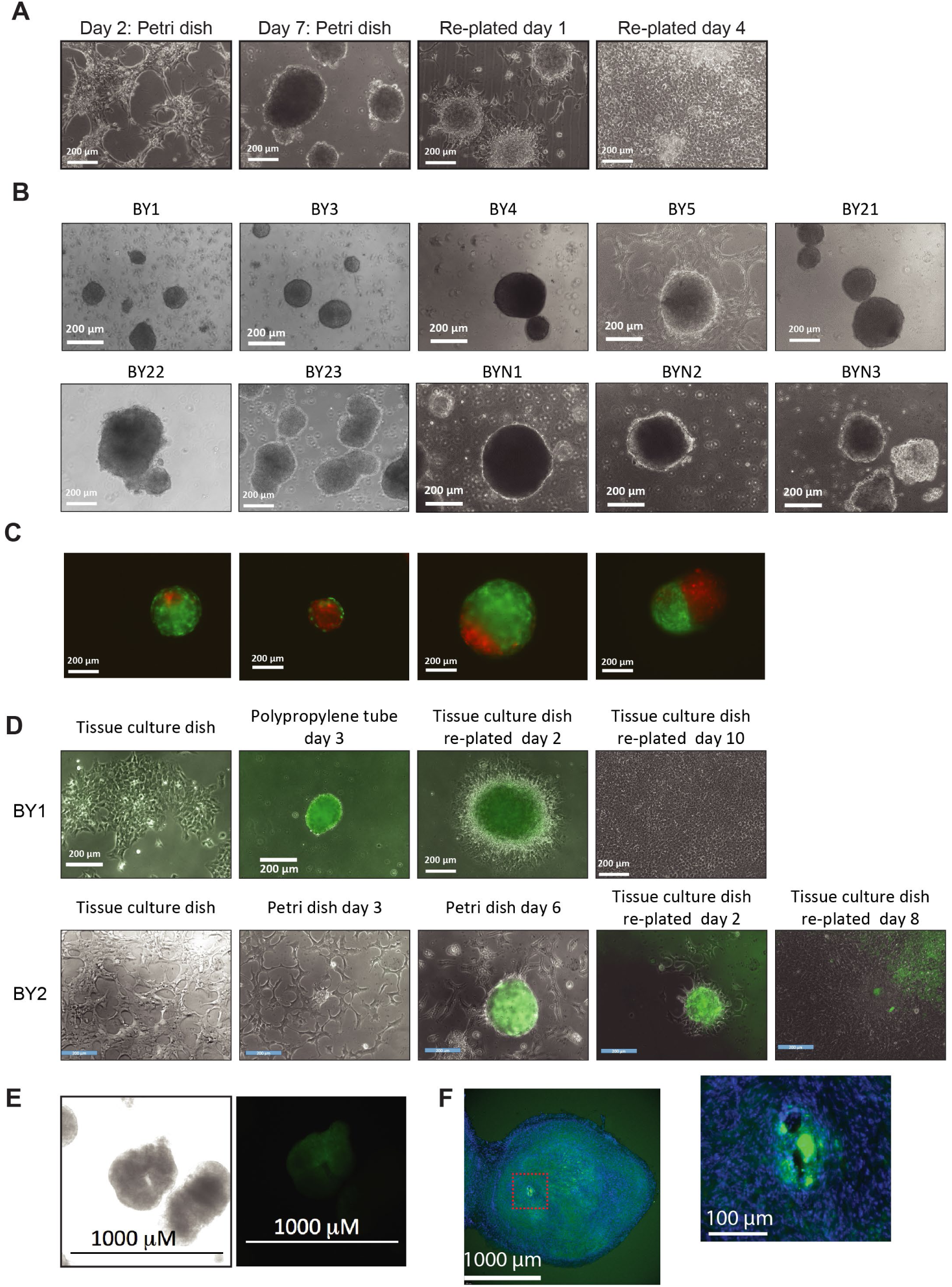
Reversible formation of tumor spheroids by HB cell lines. (A). Monolayer cultures of BY2 cells were trypsinized and re-plated onto plastic Petri dishes. After initially attaching as monolayers (Day 2), they gradually formed spheroids, completely detached from the plates by Day 7 and could be maintained in an attachment-free state for at least 4 weeks. They were then re-plated onto standard tissue culture plates and observed over 4 additional days during which time they resumed monolayer growth. (B). Spheroid formation by other HB cell lines. Single cell suspensions of the indicated BY and BYN cell lines were seeded onto bacterial Petri dishes as described in A or into polypropylene test tubes. All photos were obtained on day 7. (C). Spheroids can interact. BY2 cells stably expressing EGFP or DsRed were maintained separately, allowed to form spheroids and then co-cultured for 3 additional days before obtaining fluorescence micrographs. (D). Tumor cell spheroids activate an EC-specific promoter. BY1 and BY2 cells stably transfected with a Tie2-EGFP vector ^52^were initially maintained as monolayer cultures before being allowed to form spheroids and express EGFP. Upon re-plating spheroids onto standard tissue culture plates, GFP expression was rapidly lost as the cells resumed monolayer growth. The micrographs shown here are superimposed bright field and fluorescent images. (E). BY2 spheroids generated as described in D were exposed to the hypoxia-responsive dye Image-iT Green (10 μM for 2 hr). Bright field and fluorescent images were then obtained and superimposed. (F). BY1-Tie2-EGFP cells were grown as sub-cutaneous tumors in FVB/N mice. Frozen sections were then examined for the expression of EGFP and counterstained with the nuclear stain Hoechst 33342. The boxed region, an enlarged image of which is shown to the right, shows structures resembling blood vessels surrounded by EGFP+ cells.

Some tumor cells can “transdifferentiate” and express endothelial cell (EC)-specific markers when maintained under hypoxic conditions *in vitro* and during *in vivo* tumorigensis. ^52, 71, 72^ We therefore asked if the interiors of HB spheroids might present a similar hypoxic environment that was conducive to the expression of EC-specific markers. BY1 and BY2 cells were therefore stably transfected with a vector in which EGFP was regulated by the tightly controlled EC-specific angiopoietin receptor (Tie2) promoter. ^52^ During monolayer growth under normoxic conditions, few if any EGFP-expressing cells were observed, which mirrored previous findings in several human cancer cell lines (Figure 12D). ^52^ Spheroid formation, however, was accompanied by the appearance of numerous EGFP-positive cells, which were particularly prominent in the interiors and which rapidly diminished after resuming monolayer growth (Figure 12D). Staining with the hypoxia-detecting dye Image-iT Green also showed strong fluorescence within the interiors of many tumor spheroids but not of monolayer cultures (Figure 12 E and not shown). Thus, the temporal and spatial expression and maintenance of EGFP coincided with a hypoxic microenvironment. EGFP+ cells were also detected in the interiors of subcutaneous tumors originating from BY1-Tie2-EGFP cells and were particularly prominent around the peripheries of structures resembling blood vessels (Figure 12F).

## DISCUSSION

We have demonstrated here the feasibility of deriving immortalized isogenic cell lines from murine HBs bearing defined and clinically relevant oncogenic drivers. The B^del90^ and N^L30P^ mutants utilized for these studies were originally identified in patient tumors and were previously used to drive experimental murine HBs that bear many features of their human counterparts. ^19–21, 73^ Functionally similar mutations of B or amplification of the WT *NFE2L2* gene are also encountered in a majority of human HBs. ^1, 5, 7, 10, 15, 16, 19–21^ Although the Y^S127A^ mutant is not naturally occurring, its behavior nonetheless mimics that of HB-associated WT Y, which, in the majority of human HBs, localizes to the nucleus in a manner that emancipates it from Hippo signaling dependency and reflects a more upstream but ill-defined dysregulation of the pathway. ^19–21^ The ease with which the cell lines described here can be established from primary tumors that were previously resistant to immortalization should allow this approach to be extended to murine HBs generated by other oncogene combinations. This approach could eventually provide a comprehensive collection of cell lines that, in addition to being isogenic, are representative of any desired human HB subtype. ^7, 10, 21, 74^ Given the acknowledged paucity of human HB cell lines, it will be important in future work to determine whether the approach described here can be successfully applied to primary human HBs, thus addressing a logical next step. ^28^

Key to the high efficiency with which HB lines could be derived was the Crispr-mediated inactivation of the *Cdkn2a* locus that encodes both p16^INK4A^ and p19^ARF^.^38–40, 49, 63^ We have reported that the enforced over-expression of p19^ARF^ completely abrogates *in vivo* BY-driven tumorigenesis and similar outcomes were obtained in the current work when p16^INK4A^ was over-expressed both *in vivo* and *in vitro* (Figure 8 D and E and data not shown). ^43^ The finding that all experimental murine HBs and a significant subset of primary human HBs up-regulate p16^INK4A^ and p14/19^ARF^ and that their downstream Rb and TP53 signaling pathways remain intact (Figure 2) led us to speculate that these responses represent normal but futile attempts to limit tumor growth via the induction of senescence, apoptosis and cell cycle arrest. We further hypothesized that while the Rb and TP53 signaling pathways can be overwhelmed by the potent oncogenic signals that drive *in vivo* tumor growth, they remain sufficiently robust to block subsequent *in vitro* immortalization. Inactivating *Cdkn2a*’s exon 2, or more narrowly, the p16^INK4A^-encoding exon 1α, was sufficient to overcome the barriers imposed by these processes, thereby leading to more rapid tumor growth and *in vitro* establishment (Figure 8A). The fact that tumor cell growth could be inhibited both *in vivo* and *in vitro* by enforcing WT p16^INK4A^ or p19^ARF^ over-expression provides additional evidence that these TSs’ signaling pathways remain intact but that only their activation above a certain threshold can completely override the potent oncogenic signaling of B, Y and N (Figure 8 D and E). The fact that *in vivo* tumorigenesis was accelerated by Crispr-mediated targeting of the *Cdkn2a* locus supports this notion as does the fact that the expression of WT p16^INK4A^ and p14^ARF^ expression are found in a number of human cancers where they not infrequently are associated with longer survival (Figure 3). ^42, 51, 75, 76^

The known link between *Cdkn2a* inactivation and cellular immortalization provided our initial motivation for evaluating this locus in the context of HB pathogenesis and the *in vitro* establishment of permanent cell lines. ^38–40, 46, 48, 49^ Hyper-methylation of *Cdkn2a’s* p16^INK4A^-specific exon 1α promoter and down-regulation of p16^INK4A^ expression have been reported in approximately 50% of human HBs and a wider inactivation of the locus that includes p14^ARF^ could well account for the virtual absence of both transcripts in the primary HBs we profiled (Figure 2B & C). ^44, 77^ Thus, the cell lines that we describe here may, at least at the molecular level, be more representative of this human subset and functionally quite similar if not identical.

Given the imprecise nature of the Crispr/Cas9-mediated mutational targeting used here, its error-prone repair via non-homologous end joining ^78, 79^ and the fact that as many as 4 different gRNAs were used concurrently to facilitate the locus’ inactivation (Table 2), we anticipated that few if any HBs would express p16^INK4A^ or p19^ARF^. Thus, many of the mutations identified in both P0 and immortalized populations were comprised of large deletions or mutations that likely lacked any coding capacity (Table 4 and Supplementary File 1). Unexpectedly however, many transcripts were predicted to encode in-frame p16^INK4A^ and/or p19^ARF^ deletions or fusions and appeared to be either functionally neutral or associated with a growth advantage given their maintenance or further selection in immortalized cell lines (Figure 8C and Figure 9A). Similar fusion proteins have been predicted to occur frequently in melanomas and one that retained both its p16^INK4A^-like ability to interact with Cdk4 and its p19^ARF^-like nucleolar localization has been described in the germ-line of an individual from a melanoma-prone family. ^74, 80^ Despite this seemingly positive selection, however, all fusion proteins examined still exhibited variable levels of growth suppression of BY21 cells following their stable over-expression although this was often less extensive that it was for WT p19^ARF^ protein (Figure 10). We believe that these seemingly contradictory findings of positive selection for these mutations despite their growth suppressive effects can be explained in much the same way that WT p16^INK4^ and p19^ARF^ can be expressed by primary tumors (Figure 8B). In immortalized cells, endogenous levels of these fusion proteins retain growth suppressive functions that are weaker than those of the WT proteins. This allows them to be maintained or selected for over time in the face of strong growth-promoting oncogenic signals. Only high-level over-expression of these fusion proteins allows them to overcome the oncogenic signaling pathways and achieve growth suppression. This interpretation also explains why in BY and BYN tumors expressing endogenous levels of p16^INK4A^ and p19^ARF^ their signaling pathways remain intact but are unable to sufficiently engage their target genes, thus allowing their regulation to be subverted and overridden by oncogenic signals and regulated in directions opposite to those predicted (Figure 2D-G, Figure 3, 4, 5 and 7).

That the abundance of certain other p16^INK4A^ and p19^ARF^ mutants was markedly reduced during the course of *in vitro* propagation was consistent with the well-accepted idea that they served classical TS-like functions and conferred a proliferative disadvantage to rapidly growing, immortalized cell populations. ^39, 40, 49^ Noteworthy examples of these were seen with the mutants BY1-2α/β, BYN1-2αβ and BYN2-2α/β of which encoded similar appearing in-frame internal deletion mutants of p16^INK4A^ and p19^ARF^. This suggested that, like their WT counterparts in control tumors (Figure 8B) these mutants might have restricted tumor growth somewhat *in vivo* but were unable to do so under *in vitro* conditions associated with ideal environmental conditions and more rapid proliferation. In contrast 3 of the 4 mutations that were the most highly selected during *in vitro* propagation, namely BY1-1αβ, BY3-1αβ and BYN1-1αβ involved fusions and truncations, which were strikingly similar to those found in naturally-occurring human HCCs and other cancers (Figure 9B). Indeed, the p16^INK4A^-p19^ARF^ fusion proteins identified in several human HCCs, and that likely retained some TS-like functions, closely resembled several of our Crispr/Cas9-generated mutations involving residues 53-80 of p16^INK4A^ and residues 68-95 of p14^ARF^.

All but one of the 11 BY and BYN cell lines could be propagated in mice as either subcutaneous or pulmonary tumors while retaining the original crowded fetal-like histology of the primary HBs from which they originated (Figure 11). These cell lines should therefore be useful for elucidating the factors that contribute to both primary and metastatic growth and for determining how these behaviors are influenced by different combinations of the B,Y and N drivers. Given that these cell lines were all generated from FVB/N mice, they offer an additional advantage in that all tumor growth *in vivo* can be studied in immuno-competent hosts, thus providing an additional advantage over currently available human cell lines.

Spheroid formation allows tumor cells to be studied under conditions that more closely mimic those encountered in the hypoxic and/or nutrient-deprived tumor microenvironment. ^65, 67–70^ All 11 BY and BYN cell lines readily formed spheroids in response to plating on non-adherent or quasi-adherent surfaces (Figure 12 A and B). These could be maintained with high viability for over one month and readily resumed 2-D growth when re-plated onto a proper adherent surface. Because the conditions used for spheroid generation are identical in each case and the changes appear to be highly coordinated, the underlying molecular events that are responsible for these changes, and their timing, should to be readily quantifiable.

Tumor hypoxia contributes to metabolic rewiring and the development of resistance to a variety of therapeutic modalities. ^81–84^ It also drives neovascularization so as to re-establish a locally normoxic state that is supportive of survival and further tumor growth. The basis for this classically involves the recruitment of neighboring, extra-tumoral blood vessel-associated ECs. ^85^ However, it may also involve vasculogneic mimicry, whereby tumor cells assume the appearance and function of blood vessels without necessarily acquiring any other EC-like properties. ^86, 87^ Finally, it may involve “trans-differentiation”, an epithelial-to-mesenchymal-like transition whereby tumor cells acquire EC-like markers and properties while contributing to blood vessel formations. ^52, 71, 88^ The HB spheroid generation we have described appears to serve as a model for this latter activity in that it is associated with both the development of a rapidly reversible internal hypoxic environment and the appearance of tumor cells with at least some EC-like properties (Figure 12 D and E). This occurs *in vivo* as well and is particularly prominent in association with structures resembling blood vessels (Figure 12 F). The ability to control and quantify this process *in vitro* should allow for future studies aimed at cataloging a larger and more unbiased array of the EC-like changes that accompany spheroid formation and reversion as well as determining how and to what degree they are influenced by different combinations of oncogenic drivers.

In addition to being molecularly defined, several features of the cell lines described here will be of particular value for future studies. For example, because they are isogenic, their properties are likely to be independent of other minor oncogenic contributions and epigenetic variations that can nuance the behavior of primary tumors and obscure or modify the contributions of the primary drivers. ^7, 10, 15, 22, 23, 29, 89^ Their high transfection efficiencies, growth as either monolayers or spheroids and their ability to trans-differentiate into cells with EC-like properties will allow molecular studies that focus upon identifying the genes responsible for coordinating the transitions between these states (Figure 12 A, B and D). HBs also currently remain one of the few cancers that continue to be treated with very similar if not identical drug combinations irrespective of their underlying molecular profiles. ^90–94^ Our HB cell lines should therefore allow comparisons of how different driver oncoprotein combinations influence sensitivities to both traditional and new chemotherapeutics. The fact that most of the above HB cell lines can be propagated in immune-competent hosts and under conditions that resemble metastatic growth will certainly be helpful in this regard. Drug resistant cell lines will also be useful for investigating whether and how different driver oncogenes influence the attainment of this state ^2, 8, 13, 95, 96^.

## Supporting information

Supplementary File 1. Top mutated Cdkn2a locus sequences and the frequencies with which they were identified in 9 primary tumor-immortalized cell line

## Abbreviations

B: β-catenin
HB: hepatoblastoma
HCC: hepatocellular carcinoma
N: NFE2L2/NRF
WT: wild-type
Y: <u>y</u>es-<u>a</u>ssociated <u>p</u>rotein (YAP)

## Author contributions

W performed animals studies and bio-informatics analysis, interpreted results; JL acquired data and analyzed results; KC: acquired data and analyzed results; JK: acquired data; CH acquired data; SR performed histological analyses; EVP conceived and designed the study, drafted and revised the manuscript.

## Data Transparency

All data and cell lines generated from this study are freely available upon request

## SAGER Guidelines

All tumors and cell lines described in this study were generated from equal numbers of male and female mice

## Supplemental Material

**Supplementary File 1. Top mutated *Cdkn2a* locus sequences and the frequencies with which they were identified in 9 primary tumor-immortalized cell line pairs.**

## REFERENCES

1. Bell D, Ranganathan S, Tao J, et al. Novel Advances in Understanding of Molecular Pathogenesis of Hepatoblastoma: A Wnt/beta-Catenin Perspective. Gene Expr 2017;17:141–154.

2. Cao Y, Wu S, Tang H. An update on diagnosis and treatment of hepatoblastoma. Biosci Trends 2024;17:445–457.

3. Jeong SU, Kang HJ. Recent updates on the classification of hepatoblastoma according to the International Pediatric Liver Tumors Consensus. J Liver Cancer 2022;22:23–29.

4. Lopez-Terrada D, Alaggio R, de Davila MT, et al. Towards an international pediatric liver tumor consensus classification: proceedings of the Los Angeles COG liver tumors symposium. Mod Pathol 2014;27:472–91.

5. Prochownik EV. Reconciling the Biological and Transcriptional Variability of Hepatoblastoma with Its Mutational Uniformity. Cancers (Basel) 2021;13.

6. Ranganathan S, Lopez-Terrada D, Alaggio R. Hepatoblastoma and Pediatric Hepatocellular Carcinoma: An Update. Pediatr Dev Pathol 2020;23:79–95.

7. Cairo S, Armengol C, De Reynies A, et al. Hepatic stem-like phenotype and interplay of Wnt/beta-catenin and Myc signaling in aggressive childhood liver cancer. Cancer Cell 2008;14:471–84.

8. Carrillo-Reixach J, Torrens L, Simon-Coma M, et al. Epigenetic footprint enables molecular risk stratification of hepatoblastoma with clinical implications. J Hepatol 2020;73:328–341.

9. Cho SJ, Ranganathan S, Alaggio R, et al. Consensus classification of pediatric hepatocellular tumors: A report from the Children’s Hepatic tumors International Collaboration (CHIC). Pediatr Blood Cancer 2023:e30505.

10. Hooks KB, Audoux J, Fazli H, et al. New insights into diagnosis and therapeutic options for proliferative hepatoblastoma. Hepatology 2018;68:89–102.

11. Sindhi R, Rohan V, Bukowinski A, et al. Liver Transplantation for Pediatric Liver Cancer. Cancers (Basel) 2020;12.

12. Hubbard AK, Spector LG, Fortuna G, et al. Trends in International Incidence of Pediatric Cancers in Children Under 5 Years of Age: 1988-2012. JNCI Cancer Spectr 2019;3:pkz007.

13. Wu PV, Rangaswami A. Current Approaches in Hepatoblastoma-New Biological Insights to Inform Therapy. Curr Oncol Rep 2022;24:1209–1218.

14. Abraham SC, Wu TT, Klimstra DS, et al. Distinctive molecular genetic alterations in sporadic and familial adenomatous polyposis-associated pancreatoblastomas : frequent alterations in the APC/beta-catenin pathway and chromosome 11p. Am J Pathol 2001;159:1619–27.

15. Buendia MA. Genetic alterations in hepatoblastoma and hepatocellular carcinoma: common and distinctive aspects. Med Pediatr Oncol 2002;39:530–5.

16. Eichenmuller M, Trippel F, Kreuder M, et al. The genomic landscape of hepatoblastoma and their progenies with HCC-like features. J Hepatol 2014;61:1312–20.

17. Miao J, Kusafuka T, Udatsu Y, et al. Sequence variants of the Axin gene in hepatoblastoma. Hepatol Res 2003;25:174–179.

18. Li H, Wolfe A, Septer S, et al. Deregulation of Hippo kinase signalling in human hepatic malignancies. Liver Int 2012;32:38–47.

19. Tao J, Calvisi DF, Ranganathan S, et al. Activation of beta-catenin and Yap1 in human hepatoblastoma and induction of hepatocarcinogenesis in mice. Gastroenterology 2014;147:690–701.

20. Wang H, Lu J, Mandel JA, et al. Patient-Derived Mutant Forms of NFE2L2/NRF2 Drive Aggressive Murine Hepatoblastomas. Cell Mol Gastroenterol Hepatol 2021;12:199–228.

21. Zhang W, Meyfeldt J, Wang H, et al. beta-Catenin mutations as determinants of hepatoblastoma phenotypes in mice. J Biol Chem 2019;294:17524–17542.

22. Grobner SN, Worst BC, Weischenfeldt J, et al. The landscape of genomic alterations across childhood cancers. Nature 2018;555:321–327.

23. Ma X, Liu Y, Liu Y, et al. Pan-cancer genome and transcriptome analyses of 1,699 paediatric leukaemias and solid tumours. Nature 2018;555:371–376.

24. Noe O, Filipiak L, Royfman R, et al. Adenomatous polyposis coli in cancer and therapeutic implications. Oncol Rev 2021;15:534.

25. Zhong ZA, Michalski MN, Stevens PD, et al. Regulation of Wnt receptor activity: Implications for therapeutic development in colon cancer. J Biol Chem 2021;296:100782.

26. Dinarvand P, Davaro EP, Doan JV, et al. Familial Adenomatous Polyposis Syndrome: An Update and Review of Extraintestinal Manifestations. Arch Pathol Lab Med 2019;143:1382–1398.

27. Yang A, Sisson R, Gupta A, et al. Germline APC mutations in hepatoblastoma. Pediatr Blood Cancer 2018;65.

28. Rikhi RR, Spady KK, Hoffman RI, et al. Hepatoblastoma: A Need for Cell Lines and Tissue Banks to Develop Targeted Drug Therapies. Front Pediatr 2016;4:22.

29. Fang J, Singh S, Cheng C, et al. Genome-wide mapping of cancer dependency genes and genetic modifiers of chemotherapy in high-risk hepatoblastoma. Nat Commun 2023;14:4003.

30. Arzumanian VA, Kiseleva OI, Poverennaya EV. The Curious Case of the HepG2 Cell Line: 40 Years of Expertise. Int J Mol Sci 2021;22.

31. Doi I. Establishment of a cell line and its clonal sublines from a patient with hepatoblastoma. Gan 1976;67:1–10.

32. Pietsch T, Fonatsch C, Albrecht S, et al. Characterization of the continuous cell line HepT1 derived from a human hepatoblastoma. Lab Invest 1996;74:809–18.

33. Aden DP, Fogel A, Plotkin S, et al. Controlled synthesis of HBsAg in a differentiated human liver carcinoma-derived cell line. Nature 1979;282:615–6.

34. Lopez-Terrada D, Cheung SW, Finegold MJ, et al. Hep G2 is a hepatoblastoma-derived cell line. Hum Pathol 2009;40:1512–5.

35. Dolezal JM, Wang H, Kulkarni S, et al. Sequential adaptive changes in a c-Myc-driven model of hepatocellular carcinoma. J Biol Chem 2017;292:10068–10086.

36. Collins CJ, Sedivy JM. Involvement of the INK4a/Arf gene locus in senescence. Aging Cell 2003;2:145–50.

37. Fontana R, Ranieri M, La Mantia G, et al. Dual Role of the Alternative Reading Frame ARF Protein in Cancer. Biomolecules 2019;9.

38. Carnero A, Hudson JD, Price CM, et al. p16INK4A and p19ARF act in overlapping pathways in cellular immortalization. Nat Cell Biol 2000;2:148–55.

39. Sherr CJ, Bertwistle D, W Denb, et al. p53-Dependent and -independent functions of the Arf tumor suppressor. Cold Spring Harb Symp Quant Biol 2005;70:129–37.

40. Sherr CJ, McCormick F. The RB and p53 pathways in cancer. Cancer Cell 2002;2:103–12.

41. Cilluffo D, Barra V, Di Leonardo A. P14(ARF): The Absence that Makes the Difference. Genes (Basel) 2020;11.

42. Iolascon A, Giordani L, Moretti A, et al. Analysis of CDKN2A, CDKN2B, CDKN2C, and cyclin Ds gene status in hepatoblastoma. Hepatology 1998;27:989–95.

43. Kulkarni S, Dolezal JM, Wang H, et al. Ribosomopathy-like properties of murine and human cancers. PLoS One 2017;12:e0182705.

44. Shim YH, Park HJ, Choi MS, et al. Hypermethylation of the p16 gene and lack of p16 expression in hepatoblastoma. Mod Pathol 2003;16:430–6.

45. Gorgoulis V, Adams PD, Alimonti A, et al. Cellular Senescence: Defining a Path Forward. Cell 2019;179:813–827.

46. Lundberg AS, Hahn WC, Gupta P, et al. Genes involved in senescence and immortalization. Curr Opin Cell Biol 2000;12:705–9.

47. Mikula M, Fuchs E, Huber H, et al. Immortalized p19ARF null hepatocytes restore liver injury and generate hepatic progenitors after transplantation. Hepatology 2004;39:628–34.

48. Odell A, Askham J, Whibley C, et al. How to become immortal: let MEFs count the ways. Aging (Albany NY) 2010;2:160–5.

49. Sharpless NE, Sherr CJ. Forging a signature of in vivo senescence. Nat Rev Cancer 2015;15:397–408.

50. Zindy F, Eischen CM, Randle DH, et al. Myc signaling via the ARF tumor suppressor regulates p53-dependent apoptosis and immortalization. Genes Dev 1998;12:2424–33.

51. Witkiewicz AK, Knudsen KE, Dicker AP, et al. The meaning of p16(ink4a) expression in tumors: functional significance, clinical associations and future developments. Cell Cycle 2011;10:2497–503.

52. Elster JD, McGuire TF, Lu J, et al. Rapid in vitro derivation of endothelium directly from human cancer cells. PLoS One 2013;8:e77675.

53. Concordet JP, Haeussler M. CRISPOR: intuitive guide selection for CRISPR/Cas9 genome editing experiments and screens. Nucleic Acids Res 2018;46:W242–W245.

54. Ewels PA, Peltzer A, Fillinger S, et al. The nf-core framework for community-curated bioinformatics pipelines. Nat Biotechnol 2020;38:276–278.

55. Goldman MJ, Craft B, Hastie M, et al. Visualizing and interpreting cancer genomics data via the Xena platform. Nat Biotechnol 2020;38:675–678.

56. Hiebert SW, Chellappan SP, Horowitz JM, et al. The interaction of RB with E2F coincides with an inhibition of the transcriptional activity of E2F. Genes Dev 1992;6:177–85.

57. Timmers C, Sharma N, Opavsky R, et al. E2f1, E2f2, and E2f3 control E2F target expression and cellular proliferation via a p53-dependent negative feedback loop. Mol Cell Biol 2007;27:65–78.

58. Thut CJ, Goodrich JA, Tjian R. Repression of p53-mediated transcription by MDM2: a dual mechanism. Genes Dev 1997;11:1974–86.

59. Al-Mohanna MA, Manogaran PS, Al-Mukhalafi Z, et al. The tumor suppressor p16(INK4a) gene is a regulator of apoptosis induced by ultraviolet light and cisplatin. Oncogene 2004;23:201–12.

60. Bardeesy N, Aguirre AJ, Chu GC, et al. Both p16(Ink4a) and the p19(Arf)-p53 pathway constrain progression of pancreatic adenocarcinoma in the mouse. Proc Natl Acad Sci U S A 2006;103:5947–52.

61. Ginsberg D. E2F1 pathways to apoptosis. FEBS Lett 2002;529:122–5.

62. Mijit M, Caracciolo V, Melillo A, et al. Role of p53 in the Regulation of Cellular Senescence. Biomolecules 2020;10.

63. Kamijo T, Zindy F, Roussel MF, et al. Tumor suppression at the mouse INK4a locus mediated by the alternative reading frame product p19ARF. Cell 1997;91:649–59.

64. Young NP, Jacks T. Tissue-specific p19Arf regulation dictates the response to oncogenic K-ras. Proc Natl Acad Sci U S A 2010;107:10184–9.

65. El Harane S, Zidi B, El Harane N, et al. Cancer Spheroids and Organoids as Novel Tools for Research and Therapy: State of the Art and Challenges to Guide Precision Medicine. Cells 2023;12.

66. Manduca N, Maccafeo E, De Maria R, et al. 3D cancer models: One step closer to in vitro human studies. Front Immunol 2023;14:1175503.

67. Veninga V, Voest EE. Tumor organoids: Opportunities and challenges to guide precision medicine. Cancer Cell 2021;39:1190–1201.

68. Lv J, Du X, Wang M, et al. Construction of tumor organoids and their application to cancer research and therapy. Theranostics 2024;14:1101–1125.

69. Saltsman JA, Hammond WJ, Narayan NJC, et al. A Human Organoid Model of Aggressive Hepatoblastoma for Disease Modeling and Drug Testing. Cancers (Basel) 2020;12.

70. Ishiguro T, Ohata H, Sato A, et al. Tumor-derived spheroids: Relevance to cancer stem cells and clinical applications. Cancer Sci 2017;108:283–289.

71. Sajithlal GB, McGuire TF, Lu J, et al. Endothelial-like cells derived directly from human tumor xenografts. Int J Cancer 2010;127:2268–78.

72. Testa U, Pelosi E, Castelli G. Endothelial Progenitors in the Tumor Microenvironment. Adv Exp Med Biol 2020;1263:85–115.

73. Sylvester KG, Colnot S. Hippo/YAP, beta-catenin, and the cancer cell: a “menage a trois” in hepatoblastoma. Gastroenterology 2014;147:562–5.

74. Fung DC, Holland EA, Becker TM, et al. eMelanoBase: an online locus-specific variant database for familial melanoma. Hum Mutat 2003;21:2–7.

75. Inoue K, Fry EA. Aberrant Expression of p14(ARF) in Human Cancers: A New Biomarker? Tumor Microenviron 2018;1:37–44.

76. Inoue K, Fry EA. Aberrant expression of p16(INK4a) in human cancers - a new biomarker? Cancer Rep Rev 2018;2.

77. Pineau P, Marchio A, Nagamori S, et al. Homozygous deletion scanning in hepatobiliary tumor cell lines reveals alternative pathways for liver carcinogenesis. Hepatology 2003;37:852–61.

78. Li T, Yang Y, Qi H, et al. CRISPR/Cas9 therapeutics: progress and prospects. Signal Transduct Target Ther 2023;8:36.

79. Sansbury BM, Hewes AM, Kmiec EB. Understanding the diversity of genetic outcomes from CRISPR-Cas generated homology-directed repair. Commun Biol 2019;2:458.

80. Brookes S, Rowe J, Ruas M, et al. INK4a-deficient human diploid fibroblasts are resistant to RAS-induced senescence. EMBO J 2002;21:2936–45.

81. Chen Z, Han F, Du Y, et al. Hypoxic microenvironment in cancer: molecular mechanisms and therapeutic interventions. Signal Transduct Target Ther 2023;8:70.

82. Kopecka J, Salaroglio IC, Perez-Ruiz E, et al. Hypoxia as a driver of resistance to immunotherapy. Drug Resist Updat 2021;59:100787.

83. Lyssiotis CA, Kimmelman AC. Metabolic Interactions in the Tumor Microenvironment. Trends Cell Biol 2017;27:863–875.

84. Sorensen BS, Horsman MR. Tumor Hypoxia: Impact on Radiation Therapy and Molecular Pathways. Front Oncol 2020;10:562.

85. Perez-Gutierrez L, Ferrara N. Biology and therapeutic targeting of vascular endothelial growth factor A. Nat Rev Mol Cell Biol 2023;24:816–834.

86. Maddison K, Bowden NA, Graves MC, et al. Characteristics of vasculogenic mimicry and tumour to endothelial transdifferentiation in human glioblastoma: a systematic review. BMC Cancer 2023;23:185.

87. Wei X, Chen Y, Jiang X, et al. Mechanisms of vasculogenic mimicry in hypoxic tumor microenvironments. Mol Cancer 2021;20:7.

88. He H, Niu CS, Li MW. Correlation between glioblastoma stem-like cells and tumor vascularization. Oncol Rep 2012;27:45–50.

89. Nagae G, Yamamoto S, Fujita M, et al. Genetic and epigenetic basis of hepatoblastoma diversity. Nat Commun 2021;12:5423.

90. Brock PR, Maibach R, Childs M, et al. Sodium Thiosulfate for Protection from Cisplatin-Induced Hearing Loss. N Engl J Med 2018;378:2376–2385.

91. Czauderna P. Hepatoblastoma throughout SIOPEL trials - clinical lessons learnt. Front Biosci (Elite Ed) 2012;4:470–9.

92. Katzenstein HM, Langham MR, Malogolowkin MH, et al. Minimal adjuvant chemotherapy for children with hepatoblastoma resected at diagnosis (AHEP0731): a Children’s Oncology Group, multicentre, phase 3 trial. Lancet Oncol 2019;20:719–727.

93. Perilongo G, Maibach R, Shafford E, et al. Cisplatin versus cisplatin plus doxorubicin for standard-risk hepatoblastoma. N Engl J Med 2009;361:1662–70.

94. Zsiros J, Brugieres L, Brock P, et al. Dose-dense cisplatin-based chemotherapy and surgery for children with high-risk hepatoblastoma (SIOPEL-4): a prospective, single-arm, feasibility study. Lancet Oncol 2013;14:834–42.

95. Chow EK, Fan LL, Chen X, et al. Oncogene-specific formation of chemoresistant murine hepatic cancer stem cells. Hepatology 2012;56:1331–41.

96. Hirsch TZ, Pilet J, Morcrette G, et al. Integrated Genomic Analysis Identifies Driver Genes and Cisplatin-Resistant Progenitor Phenotype in Pediatric Liver Cancer. Cancer Discov 2021;11:2524–2543.

